# Automated identification of individual birds by song enables multi-year recapture from passive acoustic monitoring data

**DOI:** 10.1101/2025.06.26.661638

**Authors:** Sam Lapp, R. Patrick Lyon, Scott J. Wilson, Tessa A. Rhinehart, Chapin Czarnecki, Lauren Chronister, Cameron J. Fiss, Jeffery L. Larkin, Erin Bayne, Justin Kitzes

**Affiliations:** University of Pittsburgh; University of Alberta; Indiana University of Pennsylvania; American Bird Conservancy

## Abstract

Autonomous sensors and machine learning are transforming ecology by enabling large-scale observation of organisms and ecosystems. However, sensor data collected by camera traps, acoustic recorders, and satellites are typically used only to produce detections of unidentified individuals. Tracking individuals over time to study movement, survival, and behavior continues to require invasive and high-effort capture and marking techniques. Here, we introduce an automated, general approach that identifies individual animals from passive acoustic recordings based on individually distinctive vocalizations. Unlike previous approaches, ours can identify individuals in passive acoustic recordings without previously labeled examples of their vocalizations. We apply our approach to a model songbird species (Ovenbird, *Seiurus aurocapilla*), estimating abundance and annual survival across 126 locations and four years. Our approach identifies individuals with 96% accuracy. We find high Ovenbird apparent annual survival (0.70) and acoustic recapture probability (0.89) across 405 individuals. Our approach can readily be applied to other species with individually distinctive vocalizations using open-source Python implementations. Automated individual identification will broadly unlock the ability to passively recapture individual animals at the massive scale of autonomous sensing, supporting the study of population trajectories and informing proactive ecosystem management to prevent biodiversity loss.

## Introduction

In the face of global biodiversity loss, ecologists are developing innovative techniques for observing and studying organisms at the scale of populations and ecosystems. Though traditional ecological sampling protocols based on expert in-person surveys can rarely be implemented at such scales, emerging remote-sensing techniques in ecology have enabled landscape- and continental-scale monitoring. In particular, passive acoustic monitoring enables large-scale and cost-effective biodiversity monitoring of vocal taxa such as birds, bats, cetaceans, primates, and insects by recording acoustic communication signals produced by these animals^1,2^ and is being widely adopted for population monitoring programs^3–5^. Thanks to recent advances in machine learning, researchers can now develop automated tools for recognizing species-specific vocalizations^6^, enabling the efficiently generation of species presence-and-absence records at landscape scales.

Species identity, however, represents just one facet of the rich information encoded in vocalizations and captured through acoustic sensors. For example, many species of mammals, birds, and amphibians are capable of recognizing conspecific *individuals* by their vocalizations^7,8^. Individual vocal recognition (also called acoustic individual identification^9^) relies on an ‘acoustic signature’, an attribute of a vocalization that is consistently stable within an individual and unique for each individual, which are widespread in vocal animals^8,9^.

Acoustic identification of individuals provides an opportunity to passively collect acoustic recapture histories of individual animals (often termed “marked” data) without physical animal capture or marking^10^. Marked data are a critical component of many studies of demographic rates, movement, and population change, and collecting such data has long been one of the most effort-intensive approaches to ecological research^11^. Unlike traditional means of collecting marked observations, such as banding for birds, trapping for mammals, or PIT tagging for amphibians and reptiles, acoustic recapture will be non-invasive and cost-effective, unlocking the possibility of generating individual recapture histories at unprecedented scales with both new and existing passive acoustic recordings.

Compared to species recognition, acoustic individual identification in passive acoustic monitoring datasets carries a unique analytical challenge: when applying an automated individual vocal identification algorithm to large-scale passive acoustic monitoring data from hundreds of locations or long time periods, the set of individuals is not known *a priori*, and their vocalizations have likely not previously been recorded. This type of problem is known as ‘category discovery’ in machine learning because the algorithm must discover and distinguish individuals (categories) in a set of samples without any prior labeled examples of those individuals. Previous studies have performed automated individual vocal identification on a known set of individuals^10,12–14^ (discussed in SI I) or performed individual category discovery using targeted focal recordings with high signal-to-noise ratios^15–19^ (also discussed in SI I).

However, scaling these approaches from tens to hundreds or thousands of individuals is typically not feasible, and to date no applications of category discovery of individuals in passive acoustic monitoring data have been published (but see ^20^). Here, we describe a method for generating individual acoustic recapture histories from passive acoustic recordings. Our method uses unlabeled detections of a target species from an automated species classifier as training data and can be applied to typical passive acoustic monitoring datasets without requiring annotation of individuals for training. We demonstrate the effectiveness of our approach for a model songbird species, Ovenbird (*Seiurus aurocapilla*), and compare the accuracy of our approach to existing alternatives.

In summary, our approach has six steps (Figure 1):

1. Collect passive acoustic monitoring data at numerous sampling locations
2. Detect songs of the focal species using an automated species classifier
3. Train a feature extractor to find discriminative acoustic features in detected songs Training data: automated detections from Step 2, without annotations Training approach: use recording location as a pseudo-label for individual identity
4. Generate feature vectors (embeddings) for all detected songs using the feature extractor
5. Perform individual category discovery by clustering the feature vectors
6. Review clusters to create verified individual recapture histories for ecological modeling

**Figure 1:**
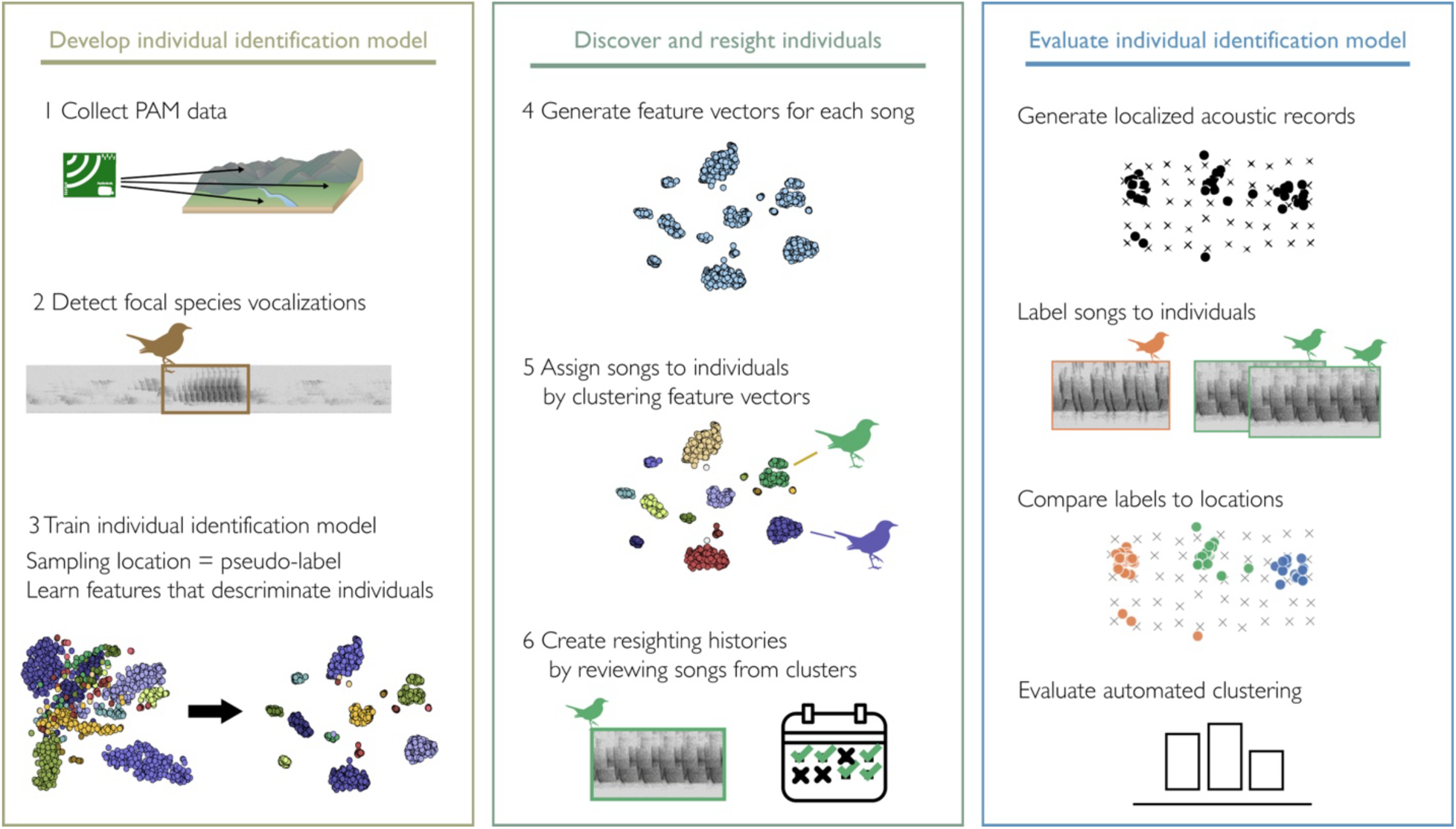
Conceptual overview for developing, applying, and evaluating automated acoustic individual identification. Though we use acoustic localization for creating evaluation data, capturing recordings of known individuals is a viable alternative.

We use the wood-warbler (*Parulidae*) species Ovenbird as a model species. Individuals of this species produce an individually distinctive primary song that is stable across years and can be differentiated by inspecting the spectrograms of song recordings^21–23^ (Figures 2, S1). During the breeding season (May-July), most male Ovenbirds defend a territory^24^ by singing their primary song^22,23^. Ovenbirds exhibit high site fidelity across years, typically establishing a territory overlapping with that of the previous year^11,25–27^. We provide a discussion of relevant Ovenbird life history in SI II.

**Figure 2:**
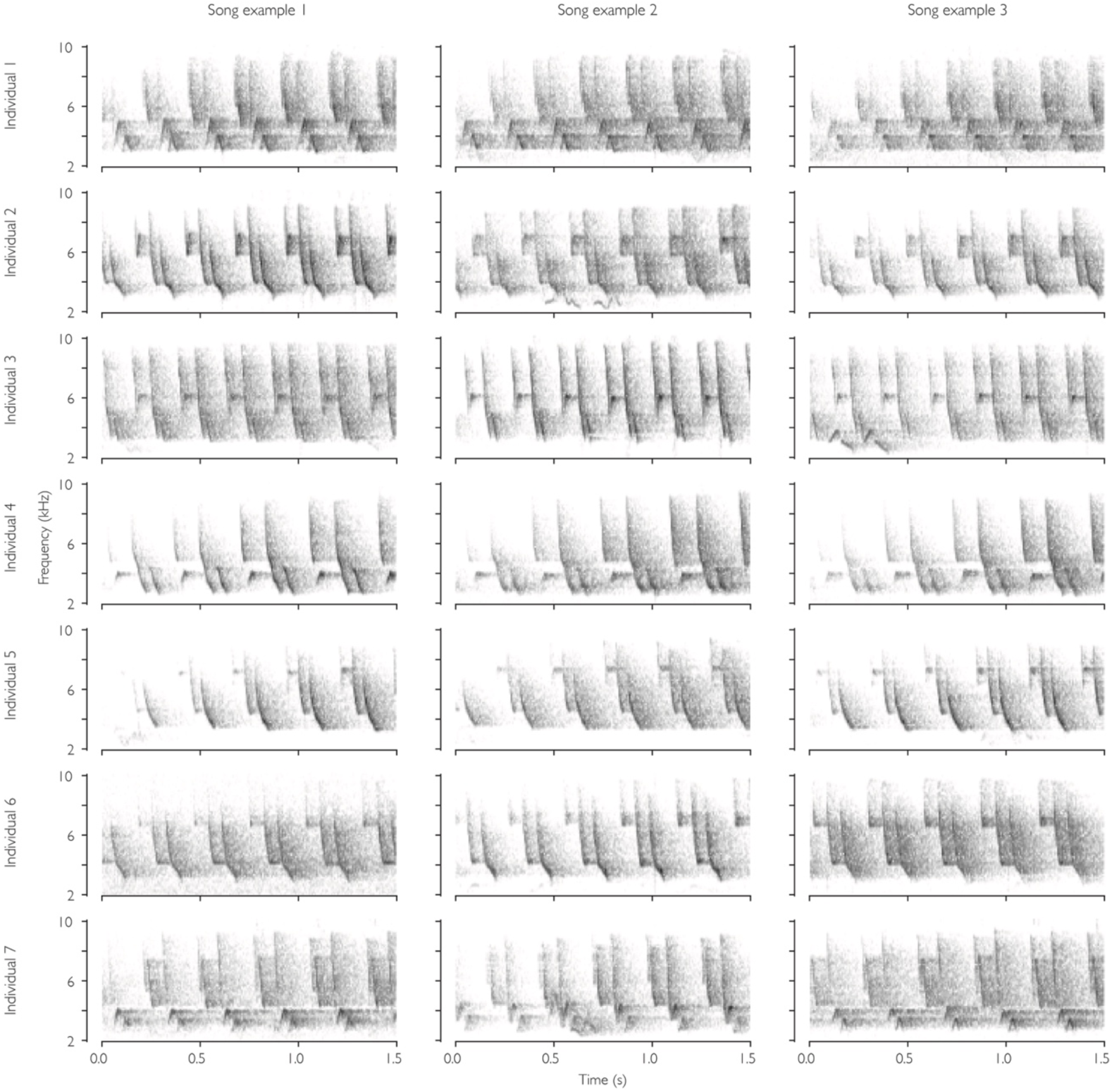
The primary songs of Ovenbirds contain stable and unique acoustic signatures. Each row shows spectrograms for three songs from an individual Ovenbird in the Localization Dataset from Alberta, Canada. Songs consist of a repeated phrase of 3-6 notes, and individuals differ in the pattern, spacing, pitch, and frequency modulation of the notes. For instance, the song of Individual 1 has a different pattern of notes than Individual 2, while the songs of Individuals 5 and 6 have similar 3-note phrases but differ in the spacing between notes. Expert observers can perceive large differences by ear in the field, while more subtle differences can only be perceived using slowed-down recordings or spectrograms. Log-valued, linear-frequency spectrograms were created from normalized 1.5-second audio clips using a 512-sample Hanning window with a step size of 256 samples and show from −55 (white) to −10 (black) dBFS using OpenSoundscape.

Ovenbirds’ primary song consists of a rapidly repeated phrase of 3-6 individual notes (Figure 2). Though many songs seem indistinguishable when heard in real time, individual Ovenbirds can consistently be identified by inspecting the acoustic features on a spectrogram^22,23^ or listening to recordings played back at reduced speed. Individuals differ in the pattern, spacing, pitch, and frequency modulation of notes that compose the song, and reproduce their song with remarkable consistency (Figure 2). Individual acoustic signatures are stable across four years, in which they return to the same location after migrating to and from their overwintering range (see Figure S1 for examples from ten randomly selected individuals across 4 years). Because Ovenbird songs are unique and stable over time, and individuals sing only one song^22^, the song can be used as an acoustic signature to distinguish and re-identify individuals in passive acoustic monitoring datasets.

We first rigorously evaluate the accuracy of our automated acoustic individual identification approach on an evaluation dataset of human-labeled songs. This dataset was annotated independently by two experts and corroborated by acoustic localization of the same songs to non-overlapping spatial territories. We demonstrate that using existing species classification models as feature extractors is not sufficient for individual category discovery, while our approach of training a supervised classifier using location as a pseudo-label is effective. We perform experiments on various parameterizations of the automated identification approach to find an optimal workflow.

We then use automated acoustic individual identification to generate individual recapture histories for 405 Ovenbirds across a four-year passive acoustic monitoring dataset, from which we measure abundance and annual survival. We provide Python implementations of our methods and open-source datasets of individually identified songs to support further development of automated individual identification tools for passive acoustic monitoring.

## Results

### Developing an annotated set of localized acoustic recordings

We performed acoustic localization of Ovenbird songs across 13 recording arrays, then annotated the localized songs with individual labels. This produced an annotated dataset with 45 individual Ovenbirds (1-8 per localization array), containing 3,963 audio clips from 717 spatially localized singing events (Figure 3; detailed results in SI VII). Agreement for individual song labels across two annotators was 98.8%, which is better than expert annotator agreement rates typical for avian acoustic species identification tasks (e.g., 23-85%^28^). Furthermore, labels assigned by human annotators generally corresponded to spatial clusters, which we interpret as individual birds’ territories (Figure 3). We use these labels to evaluate automated category discovery performance. Ten arrays are used as a validation set to select optimal parameters, and 3 as a held-out test set for final evaluation. None of the audio data or individuals from any localization array were included in training data for developing feature extractors.

**Figure 3:**
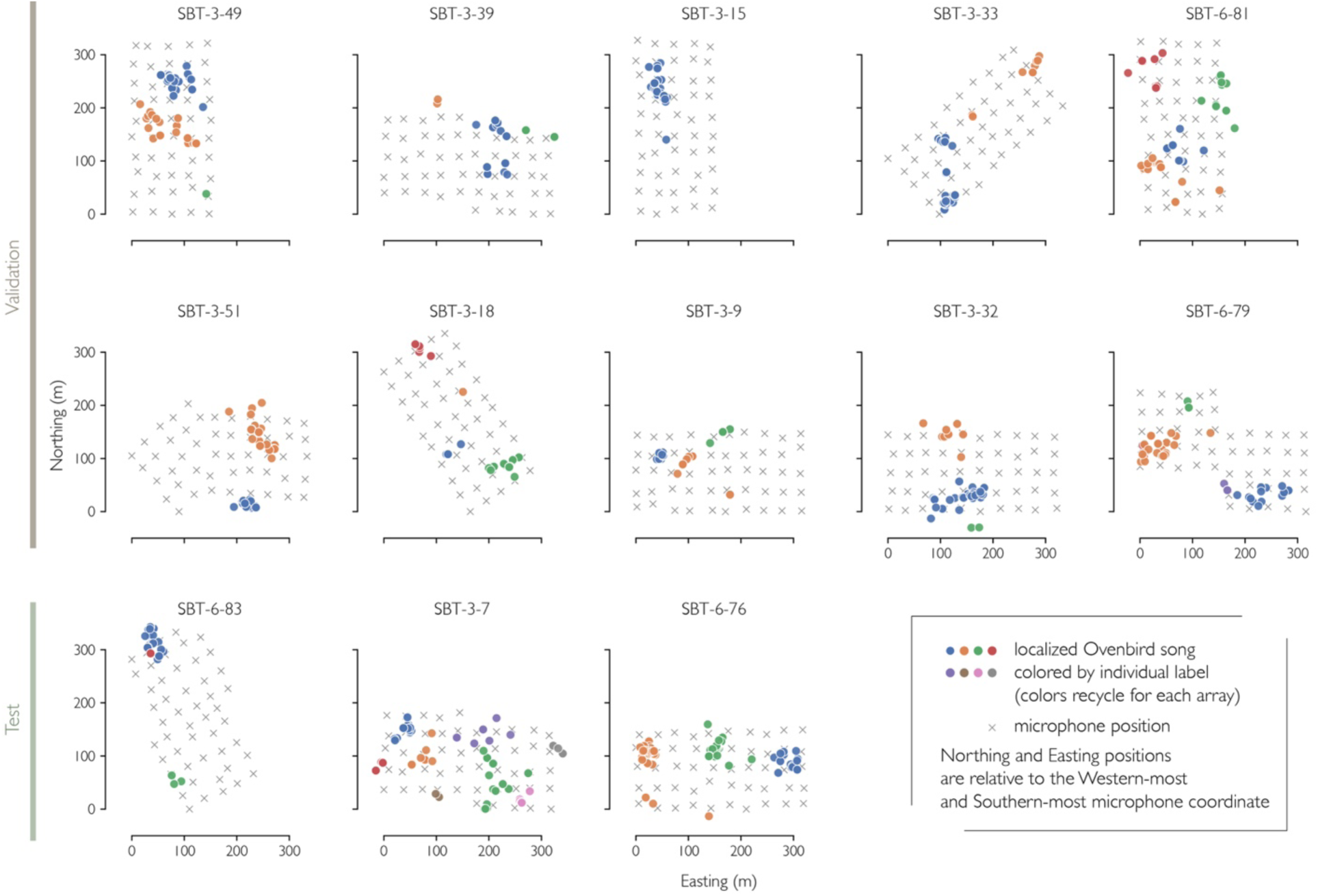
Ovenbird song positions on acoustic localization grids, colored by individual-level annotation. Interactive maps and song samples from each individual can be explored on the public web app.

We provide a public dataset containing the spatial information, individual labels, and associated audio clips for these Ovenbird songs (Environmental Data Initiative edi.2049.1), and an interactive web app for exploring audio samples and maps of individuals’ songs (https://ovenbird-id.streamlit.app/).

### Automated individual identification model

Our approach to automated individual identification involves developing a deep learning model (henceforth feature extractor) that takes a spectrogram as input and outputs a 1-dimensional feature vector. The feature extractor should extract salient acoustic features for individual identification of the target species, such that the feature vectors for songs of each individual cluster in the feature space (Figure 1 step 3).

We compared supervised and contrastive loss functions for training the deep learning feature extractor (see Methods). Supervised loss functions produced feature extractors that effectively clustered individuals’ songs, while contrastive loss functions did not (Figures S2 and S3, Table S1). Automated individual identification performance increased with the size of the training set (Figure S4a, Table S2) and with the number of unique acoustic sampling locations included in the training set (Figure S4b, Table S3). Training experiment results are discussed further in SI VIII and visualized in an online Weights and Biases report (https://bit.ly/ovenbird-training-experiments). Based on these experiments (Tables S4, S5, S6), we trained a final Ovenbird feature extractor with cross entropy loss on the entire training set. We used a ResNet18 backbone with 3-second clip inputs and included noise-reduction and overlay (weighted averaging of training samples with background clips) in audio preprocessing.

We also investigated how dimensionality reduction of features before clustering impacted performance (detailed results in SI IX). We found that reducing the dimensionality of feature vectors before performing automated clustering improved accuracy from 75% to 89% on the validation set (Table S7). We selected t-SNE^29^ reduction to 3 dimensions as the optimal approach. Reducing features to 30 dimensions with the UMAP algorithm^30^ favored in previous work^15,31^ was also effective, but performance was less consistent across runs of the stochastic algorithm and degraded when using fewer than 30 output dimensions (Figure S5).

### Assessing automated individual identification performance

The final Ovenbird feature extractor trained using our approach extracted features that resulted in accurate automated individual identification in both the validation set (accuracy 94%, ARI 0.93) and held-out test set (accuracy 0.99, ARI 0.98). This feature extractor greatly outperformed existing bird species classification models (BirdNET, HawkEars) and image classification models (ResNet18 trained on ImageNet), which were generally ineffective for automated category discovery (Tables 1, S8, Figure S6). Visualizing the individual song feature embeddings reduced to 2 dimensions with t-SNE shows the custom-trained Ovenbird feature extractor model creates dense, separate clusters for each individual (Figure 4). Features from other models still place songs from one individual close together, but do not provide enough separation to perform unsupervised clustering when labels are not known a priori (see Figure S7 for clusters assigned by HDBSCAN using each feature extractor).

**Figure 4:**
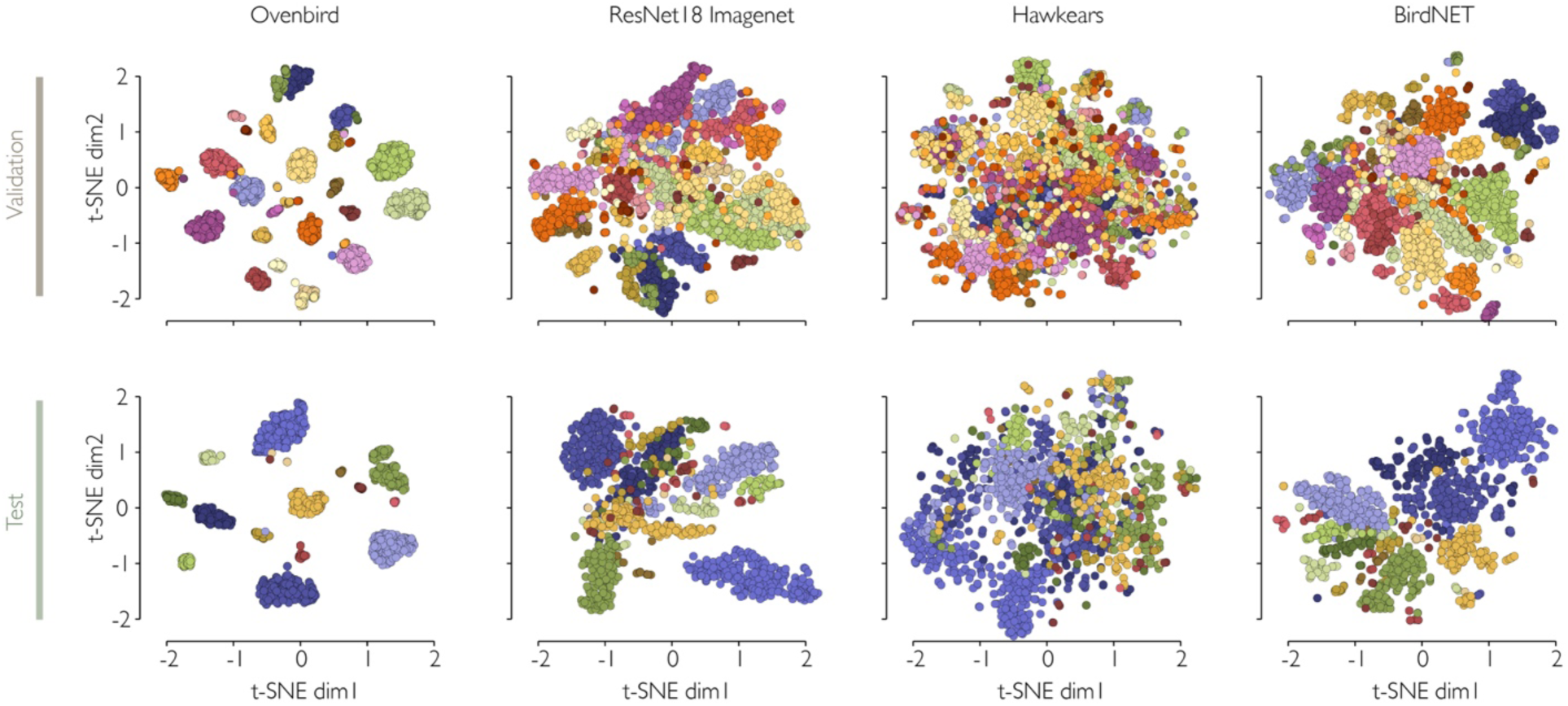
Embedding vectors for individually labeled Ovenbirds in the validation and test sets for our Ovenbird feature extractor and baseline models. Colors are the human-annotated labels. Feature vectors were generated using each of four feature extractors (columns) and reduced to 2 dimensions with t-SNE.

**Table 1:**
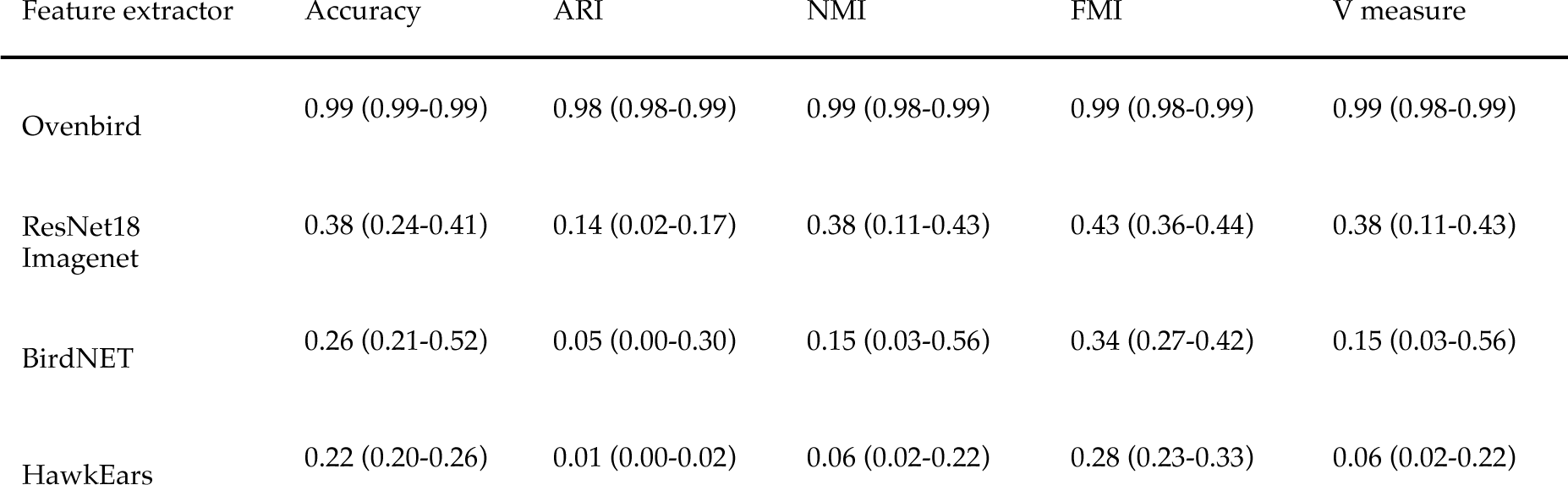
Individual identification performance on the test set when using our Ovenbird feature extractor and pre-trained baseline feature extractors. Mean and range across 30 runs are given.

| Feature extractor | Accuracy | ARI | NMI | FMI | V measure |
| --- | --- | --- | --- | --- | --- |
| Ovenbird | 0.99 (0.99-0.99) | 0.98 (0.98-0.99) | 0.99 (0.98-0.99) | 0.99 (0.98-0.99) | 0.99 (0.98-0.99) |
| ResNet18<br>Imagenet | 0.38 (0.24-0.41) | 0.14 (0.02-0.17) | 0.38 (0.11-0.43) | 0.43 (0.36-0.44) | 0.38 (0.11-0.43) |
| BirdNET | 0.26 (0.21-0.52) | 0.05 (0.00-0.30) | 0.15 (0.03-0.56) | 0.34 (0.27-0.42) | 0.15 (0.03-0.56) |
| HawkEars | 0.22 (0.20-0.26) | 0.01 (0.00-0.02) | 0.06 (0.02-0.22) | 0.28 (0.23-0.33) | 0.06 (0.02-0.22) |

For the threshold score of 1.0, the species classifier’s precision was 0.97 and recall was 0.69. After applying additional filtering to select high-quality Ovenbird song detections for individual identification (see Methods), precision was 1.00 and recall was 0.17. We found that 97% of these selected songs were assigned to a cluster during automated individual identification; thus, end-to-end recall of Ovenbird songs in the automated identification pipeline was 0.16. Low song-level recall is acceptable because individuals typically produce many detectable songs; however, future work should consider alternative approaches to our mute-and-classify approach to centering songs within clips (see detailed discussion in SI X).

We found that our individual identification model can simultaneously distinguish between the songs of hundreds of individuals (Figure 5). After preventing HDBSCAN from creating one cluster, an average of 234 clusters were detected from songs pooled across all 119 points where Ovenbird were detected, indicating that our feature extractor can discriminate between hundreds of individual Ovenbirds’ songs. Cluster purity is high, meaning that the songs are not too similar to consistently differentiate (Figure 5b).

**Figure 5:**
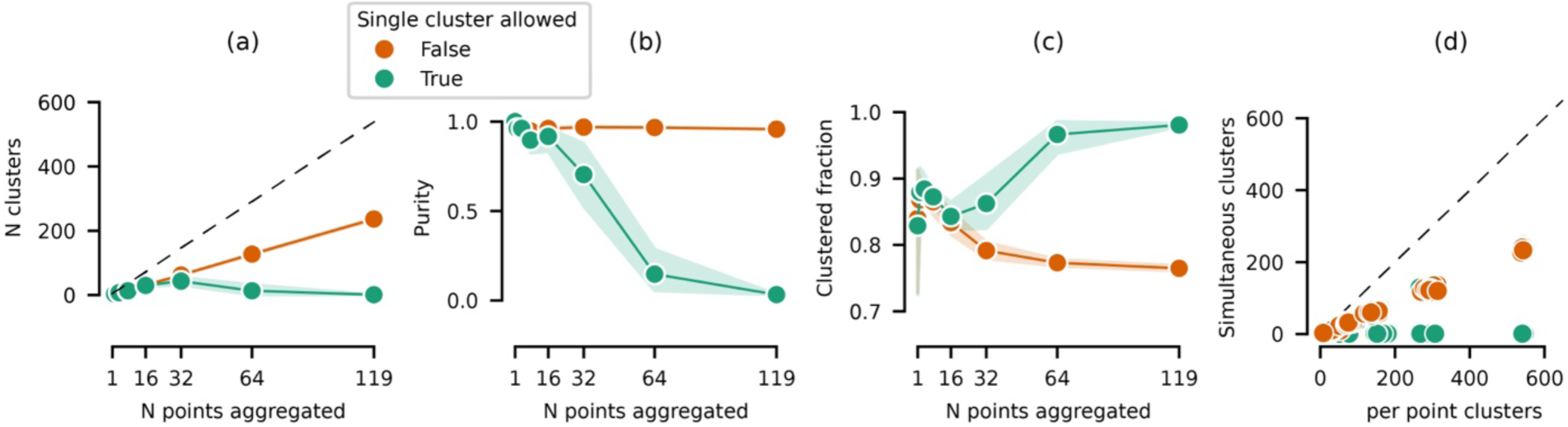
Our approach can simultaneously distinguish hundreds of individual Ovenbirds by song. When clustering songs aggregated across an increasing number of points, we plot (a) the number of clusters generated, (b) cluster purity (see definition in text), (c) the fraction of songs assigned to a cluster, and (d) the number of clusters against the number of clusters created when songs from each point are clustered separately. We show results with the HDBSCAN allow_single_cluster parameter disabled and enabled. Shaded regions show 95% confidence intervals across 20 runs.

We also assessed the accuracy of automated Ovenbird species detection with the HawkEars classifier by annotating a set of 1000 randomly selected audio clips (add dataset link once public). We found that HawkEars had excellent performance for automated detection of Ovenbird Song (AU-ROC: 0.985, AP: 0.926, details given in SI X).

### Ovenbird survival and abundance analysis

We applied our automated individual identification approach to a four-year passive acoustic monitoring dataset with 126 sampling points to estimate Ovenbird survival and abundance. Overall, our approach effectively discovered individuals in the passive acoustic monitoring dataset and frequently recaptured individuals returning to a point year after year. By applying an efficient manual review process to the results of our automated pipeline, we generated a 12-day detection history for 405 individual Ovenbirds in each of 4 years. This produced a detailed history of 19,440 individual detection/non-detection records from 9,072 hours of passive acoustic monitoring data with less than 20 hours of human review effort.

We estimated apparent annual survival of Ovenbird by fitting Cormack-Jolly-Seber (CJS) models to the four-year detection histories of 405 individuals. We found that apparent survival was 0.70 (95% credible interval 0.66-0.74) and annual detection probability was high (0.89, 95% CI 0.85-0.93, posterior distribution and summary statistics in Figure S8 and Table S9). Previous multi-year recapture studies on Ovenbirds have reported similar or lower rates of apparent survival. Vernoillet et al.^26^ found a decreasing trend of annual apparent survival from 85% in 2006 to 50% in 2014. Other studies estimate apparent survival of 61% in Missouri^27^, 62% in central Saskatchewan^32^, and 53-65% across regions in the long-term MAPS dataset^33^.

In addition to the basic CJS, we fit model variations that allowed survival to vary with year and study area, and with point-level habitat covariates associated with Ovenbird habitat selection (tree basal area, percent canopy cover, and ground slope; see Methods). Our full model estimated that apparent survival was marginally higher in the first to second year, did not differ significantly by study area, and was not explained by point-level habitat covariates (Table S10, Figure S9). See SI XI for detailed results of all models, including additional models which only considered ‘resident’ individuals detected on at least 4 days in a year.

We used generalized linear models to investigate differences in Ovenbird abundance across two study areas (State Game Lands 34 and Ohiopyle), years, and point-level habitat covariates. The full model, including basal area, percent canopy cover, and slope as predictors of abundance, was the best supported. The model without slope was nearly as well supported (delta AICc 1.0), while models removing other covariates were substantially less supported (delta AICc >=6.4, Table S11).

Average Ovenbird abundance was higher (p=6e-9) at SGL 34 (mean of 1.9-2.4 individuals per point across years) than at Ohiopyle (1.0-1.3 individuals per point across years) (Table S12). Abundance was 23% higher than 2021 in 2022, and similar to 2021 in 2023 and 2024 (Table S12). In the full model, Ovenbird abundance was positively associated with percent canopy cover (p=.006) and basal area (p=0.00006), and negatively associated with slope (p=0.08) (Table S13). Partial effects are plotted in Figure S10.

Our findings that Ovenbirds occur at higher densities in areas with higher percent canopy cover and basal area, and lower ground slope, are in line with expectations based on previous research^34^. However, these covariates did not explain apparent survival, suggesting that survival and site fidelity may be driven by different factors than habitat selection, or by habitat features we did not explore in this study.

## Discussion

We introduced an approach for performing automated acoustic individual identification of unfamiliar individuals. Notably, our approach does not involve annotation of individuals to develop training data and is able to perform category discovery of individuals, identifying novel individuals in new data rather than classifying songs to previously-known individuals. When applied to Ovenbirds, our fully approach identified unfamiliar individuals with high accuracy (validation set accuracy 94% and ARI 0.95, test set accuracy 99% and ARI 0.98) without manual review. By contrast, using pre-trained feature extractors resulted in poor performance (accurach 50%, ARI<0.22). In the absence of an automated recognizer, annotation of randomly selected Ovenbird detections from a species classifier would result in underestimates of abundance and survival (SI XII, Figure S11). Thus, using automated individual identification to acoustically recapture individuals across a large passive acoustic monitoring dataset is key for studying population trends and demographic rates.

Because we did not capture and mark individuals to create ground truthed evaluation data, we should critically consider whether our song variant labels truly correspond to individual Ovenbirds. In agreement with previous studies^22^, we found evidence that Ovenbird song variants can be reliably identified both by humans and automated procedures; that an individual is unlikely to sing more than one song variant; and that individuals’ songs are uniquely distinguishable even in a pool of over 200 individuals (detailed discussion in SI XIII). Taken together, these findings provide strong evidence that Ovenbird song variants, as labeled by human or automated approaches, correspond one-to-one with individual birds.

As with any observation process, acoustic recapture is likely to produce occasional misidentifications. We note that physical marking and resighting techniques produce valuable insights despite expert observers occasionally misidentifying individuals (e.g. misreading color bands with error rates of 0.3-6.6%^35^. We discuss how the most likely forms of acoustic misidentification would affect estimates of survival and abundance in SI XIII.

In both species classification and individual identification, manual review of automated analyses remains necessary for reliable ecological inference, as small numbers of incorrect detections can have outsized impacts on downstream ecological inference. For instance, small species classifier false positive rates can inflate occupancy estimates, while individual classifier false detections can inflate survival estimates. Rather than producing a final, error free recapture history, our approach should thus be viewed as enabling human annotators to efficiently produce accurate recapture histories that leverage the scale of passive acoustic monitoring data.

Our proposed approach to developing an automated individual identification pipeline can be readily applied to other species with stable and unique individual acoustic signatures, which are found in many taxa including mammals, birds, and amphibians^8,9^. Re-identification and counting of individuals will present additional challenges for species in which individuals sing a repertoire of multiple songs, which is the case for most songbirds^36^. In such cases, individuals’ repertoires could be delineated by identifying individually distinctive sets of songs or identifying an acoustic signature in the qualities or the sequencing of notes or phrases. More broadly, individual identification will be difficult for species that add to or modify their repertoire throughout their life, or when variation within individuals exceeds variation between individuals.

Historically, generating individual-level capture histories for birds and other taxa has required time-intensive physical capture methods that modify animal behavior and make extensive spatial replication infeasible. For example, Venouillet et al.^26^ invested 800-1600 observer hours per year to generate individual recapture data for 330 Ovenbirds through color banding and resighting at eight 25 ha plots. Such mark-recapture datasets are invaluable for understanding individual and population-level ecological processes, but cannot feasibly be expanded to cover hundreds of unique sampling points or species, and thus typically lack the statistical power needed to investigate ecological variables driving population change across communities.

Given that many researchers are already conducting passive acoustic monitoring with hundreds to thousands of recorders^5^, our approach could rapidly expand the availability of spatially replicated individual-level recapture data for a broad range of vocal species.

By achieving high spatial and temporal coverage, larger sample sizes, and higher recapture probabilities, individual acoustic monitoring will offer future opportunities for robust design surveys^37,38^ and spatial capture-recapture^39^ while increasing precision and statistical power. Passive sensor data often contains spatial and temporal autocorrelation and may warrant the development of novel statistical analyses (See SI XIV for extended discussion).

## Conclusions

Despite modern technological advances in remote sensing and artificial intelligence, contemporary animal recapture studies largely still rely on century-old capture-recapture techniques that are highly invasive and extremely time-consuming. Considering the value of individual-level recapture histories and the difficulty of obtaining them, we believe that methods for automated individual identification hold great potential for expanding the richness of ecological insights from acoustic monitoring and other passive sensing data. Automated methods capable of individual identification will produce acoustic recapture histories at the massive scales of passive sensor data, thereby contributing to our ability to study individual-level processes, investigate population trajectories, and ultimately inform proactive ecosystem management to prevent biodiversity loss.

## Acknowledgments

This material is based on work supported by the National Science Foundation under Grants 1935507, 2330423, and 2120084. This work was also supported by the Gordon and Betty Moore foundation under grant 10540. We thank The University of Pittsburgh Center for Research Computing for the resources provided. This work used the Bridges2 system at the Pittsburgh Supercomputing Center through allocation BIO240339 from the Advanced Cyberinfrastructure Coordination Ecosystem: Services & Support (ACCESS) program, which is supported by U.S. National Science Foundation grants #2138259, #2138286, #2138307, #2137603, and #2138296.

This work was also supported by the National Fish and Wildlife Foundation’s Central Appalachia Habitat Stewardship Program under grants 59680 and 66207. Collection of the Wilson and Bayne (2018) dataset was supported by a Natural Sciences and Engineering Research Council of Canada Collaborative Research and Development Grant (CRDPJ 469943-14) in conjunction with Alberta-Pacific Forest Industries, Cenovus Energy, and Conoco-Phillips Canada. Additional funding was provided by the Natural Sciences and Engineering Research Council of Canada (NSERC) CREATE and CRD programs, the Joint Oil Sands Monitoring (JOSM), the Alberta Upstream Petroleum Research Fund (AUPRF), the Canadian Oil Sands Innovation Alliance (COSIA), Environment and Climate Change Canada’s Habitat Stewardship Program (HSP), and the University of Alberta Northern Research Awards (UANRA).

## Data Availability Statement

The code associated with the methods, workflow, analysis, and figures presented in this article are available at https://github.com/sammlapp/ovenbird-individual-recognition. A model checkpoint for the Ovenbird individual feature extractor is also included in the GitHub repository. The Localization Dataset of annotated and localized Ovenbird songs is publicly available through the Environmental Data Initiative (edi.2049.1). An interactive web page visualizing maps of the labeled songs and providing audio examples with full-speed and quarter-speed playback is hosted at https://ovenbird-id.streamlit.app/. The dataset of 1000 audio clips annotated for Ovenbird song presence is available on Dryad (in prep). The Longitudinal Dataset of Ovenbird song audio clips from the 4-year passive acoustic monitoring study is available on Dryad (in prep).

## Methods

We describe an approach for developing and applying an automated procedure for individual identification of unfamiliar individuals of a focal species in passive acoustic monitoring data. Our approach uses detections of a focal species from passive acoustic monitoring data collected at multiple survey locations as training data. We do not annotate the training samples to individuals. Instead, we treat the survey location as a pseudo-label for individual identity.

Multiple individuals are likely to be detectable at a single survey location, but given sufficient spacing between survey locations, a single individual is unlikely to be detected at multiple locations. Therefore, the strategy of using survey location as a pseudo-label groups multiple individuals under a single label but does not assign different labels to songs from a single individual. We evaluate various strategies for machine learning model training given this data type.

We treat acoustic individual identification of unfamiliar individuals as a clustering problem. In contrast to a closed set classification problem, where the classes are known a priori, or an open set classification problem in which some classes are known a priori, we do not have a pre-defined set of categories (e.g., known species or individuals) to which songs should be assigned. Instead, the objective is to generate a set of clusters (categories) and assign songs to them such that two conditions are met: (1) each cluster contains only songs from one individual, and (2) all samples from one individual occur in the same cluster.

### Passive acoustic monitoring data collection

We use two existing passive acoustic monitoring datasets to develop and evaluate automated approaches to acoustic individual identification. The Longitudinal Dataset is a 4-year passive acoustic monitoring study of forest bird communities in Pennsylvania, USA^40^ from 234 survey locations across two study areas (State Game Lands 34 and Ohiopyle State Park) in Pennsylvania with a minimum of 250-meter spacing between survey locations. We use the Longitudinal Dataset as an unlabeled data source for training, for exploratory evaluation of automated individual identification behavior, and for density and survival analyses (see Case Study). The Localization Dataset contains data from 13 acoustic localization arrays in Alberta, Canada, with 50 recorders per array. Acoustic localization arrays use closely spaced (e.g., 35 m) time-synchronized recorders such that individual sound sources can be spatially localized through a post-processing procedure^1,41,42^. We use the Localization Dataset to evaluate the performance of automated individual identification methods and to validate human annotator labels of individual songs.

The Longitudinal Dataset surveyed each location each year from 2021 to 2024, from May through July. AudioMoth ARUs^43^ (v1.1, firmware 1.7.0 & 1.8.1). ARUs captured breeding bird vocal activity by recording audio continuously from 6:00 AM to 7:30 AM daily. Each ARU was configured to record with a 32 kHz sampling rate and on the medium gain setting. To minimize acoustic masking and ensure that recordings were taken omnidirectionally, ARUs were deployed on trees at breast height and a height of 1.5-2.0 meters. When possible, we selected trees <26cm in diameter at breast height. Each ARU was deployed in an anti-static waterproof bag with a desiccant pack situated behind the device and attached to a tree at each sampling point using a cable tie.

The Localization Dataset^44^ includes data from 13 acoustic localization arrays, each containing 50 recorders in rectangular layouts (Figure 3), with an average spacing between recorders of 33.9m. Each array recorded audio for 1-5 consecutive days during the breeding season (May 26 - June 27, 2016), across the peak hours of avian activity (29 minutes on, one minute off, from 05:00 to 08:30). This project used GPS-enabled Wildlife Acoustics SM3 ARUs (Wildlife Acoustics, Concord, MA, USA), where each ARU recorded audio from two locations, one from the on-board microphone and one from an external SMM-A1 microphone at a different position and connected to the ARU with a cable. Recorders used a 48 kHz sampling rate with gain set to 19.5 dB. The positions of each microphone were measured with a Hemisphere S320 survey GPS, and when the survey GPS failed to obtain GPS coordinates, the position of the ARU was determined using the on-device GPS. For further details on the collection of this dataset, see Wilson and Bayne^44^.

### Automated species detection

We use the HawkEars^45^ v0.1.0 open-source automated species classification model to detect Ovenbird songs in passive acoustic monitoring data. The species classifier produces scores representing its confidence that Ovenbird (and over 300 other vocal species) are present in consecutive 3-second audio clips. We evaluated the classifier’s performance by annotating 1000 randomly selected 3-second clips from the Localization Dataset for the presence of Ovenbird song and measuring average precision and area under the receiver-operating characteristic curve. A histogram of scores and curves of precision and recall versus score threshold is plotted in Figure S12.

In preliminary experiments with automated individual identification, we observed that feature separation between individual Ovenbird songs degrades when other sounds occur in combination with Ovenbird song in an audio clip and when the song only occurs on the edges of a clip. Therefore, we post-process species detection results in two steps. First, we filter out Ovenbird detections for which the classifier also detected other species above a threshold score. Second, we repeat the species classification on 3-second clips in which the audio outside of the central 1 second is muted and reject clips scoring below a logit score of 0.0. With this approach, we automatically filter the original detections to those where the Ovenbird song occupies the center of the audio clip and other species are not present.

### Training dataset

We created a training dataset for machine learning model development by selecting an unannotated random sample of Ovenbird songs from the Longitudinal Dataset stratified by survey location. We first used HawkEars to detect Ovenbird songs across the 4-year longitudinal dataset. We used a threshold of 1.0 on the logit score for Ovenbird detection and excluded clips if they also scored above 0.0 on the logit score for any other class (i.e., HawkEars detected another bird, mammal, or frog species in the sample). Unlike in the survival and abundance analyses (see below), for the training set, we selected audio from across all points and dates in the dataset. We dropped points with fewer than 20 Ovenbird song detections, then randomly subset to 500 detections per point. This resulted in a training set containing 94,378 3-second audio clips stratified across 126 survey points and spanning 4 years. Because HawkEars occasionally produces false-positive detections, a small fraction of these clips does not contain Ovenbird song.

We additionally selected a random sample of audio clips without Ovenbird songs as a supplementary data type for machine learning training. We randomly selected 30 3-second clips in which Ovenbird was not detected by HawkEars (logit score less than −1.0) from each recorder in the years 2022-2024.

### Evaluation dataset

We created a manually annotated dataset of Ovenbird individual songs from the Localization Dataset to evaluate the performance of automated individual identification methods.

Evaluating individual identification methods trained on the Pennsylvania training set on the Alberta validation and test sets not only ensures that the individuals in the test set do not overlap with individuals in the training set but also presents a potentially challenging domain shift. The Localization Dataset used for evaluation was recorded on different recording hardware, in a different geographic region and ecological community than the training data and contains a different population of Ovenbirds that might have different population-level acoustic characteristics.

We used automated species classification and automated acoustic localization to create a dataset of Ovenbird songs with associated spatial positions on the 13 localization arrays in the Localization Dataset. Two annotators independently inspected songs and categorized them into song variants based only on the acoustic characteristics of the song, without using the spatial position. We compared annotations across the two annotators and resolved conflicts, then compared our song variant annotations to spatial clustering of songs on the localization arrays by creating maps of songs colored by song variant. We provide a public interactive web app to explore these spatially mapped songs with individual annotations (https://ovenbird-id.streamlit.app/). Because song variants corresponded to spatially clustered songs (Figure 3), which we interpret as individual Ovenbird territories, we consider each song variant to represent an individual Ovenbird and use the annotated songs to evaluate the performance of automated acoustic individual identification methods.

### Automated species detection

We first applied the HawkEars species classifier to the Localization Dataset to detect Ovenbird songs. When selecting clips for acoustic localization, achieving moderate to high recall is important because several detections of the same singing event across spatially adjacent recorders are required to localize the song. We selected a threshold logit score of −1 for Ovenbird song because this threshold achieved high precision (91%) and recall (83%) on 3-second clips, and recall dropped quickly at higher thresholds (Figure S12). Because detections would be reviewed after localization (see below), we did not manually review automated detections at this stage.

### Automated acoustic localization

We next estimated the 2-dimensional horizontal position of Ovenbird songs detected by the automated classifier using the localization module of the open-source Python package OpenSoundscape (Lapp et al, Freeland Haynes in prep). Briefly, acoustic localization involves grouping simultaneous detections of Ovenbird songs across neighboring microphones, using generalized cross correlation to determine the relative arrival time of the song at each microphone, and computing the estimated position of the sound source based on the sound’s relative time of arrival at each of the microphone positions (see Freeland-Haynes et al for details on the procedure and the Opensoundscape implementation). We required that 5 microphones within a radius of 60 meters detect an Ovenbird song to attempt localization, and discard localized sounds with low cross-correlation scores (less than 0.01) or high residual error in the time delays versus the estimated position (root-mean-square residual error > 5 m). See Freeland-Haynes et al (in prep) and the Opensoundscape documentation for a discussion of these parameters. We further post-process the localized detections using a novel automated check of correct cross-correlation alignment (details in SI IV).

Because the acoustic localization process in Opensoundscape produces redundant position estimates of the same singing event based on different subsets of acoustic recorders, we removed duplicated singing events that occurred at the same time and within 10 meters of each other, retaining only the singing event with the lowest residual error in the location estimate.

This process produced a set of automatically detected and localized Ovenbird singing events for each acoustic localization array. Each singing event includes a set of audio clips from five or more microphones that simultaneously recorded the singing bird.

### Annotation

We manually reviewed Ovenbird singing events from one localization grid at a time to create a labeled dataset of individual Ovenbird songs for model evaluation. We also use the manual annotation of songs from the localization grid to evaluate the annotators’ ability to consistently identify individual Ovenbirds by comparing annotations from two annotators to each other and to the spatial positions of the songs.

We selected three of the 13 localization arrays as a test set and used the remaining 10 as a validation set. For the validation set, we randomly selected 50 singing events per array to annotate. For the test set, we randomly selected 100 singing events per array to annotate, or all events if there were fewer than 100. We used model performance on the validation set to tune hyperparameters of the machine learning models and the dimensionality reduction and clustering steps (see below), and the test set as an additional test of model performance.

Two reviewers independently assigned individual song variant labels to each song. Recall that each singing event from a localization grid contains a set of audio clips from microphones that simultaneously recorded the song from different locations. We reviewed the audio clip from the microphone closest to the sound source position, as recordings from shorter distances retain more of the acoustic details that differentiate individuals’ song variants. The annotator visually inspected spectrograms and listened to audio playback at one-quarter speed, which facilitated the perception of fine-scale temporal features not easily perceived in full-speed playback. The annotator did not know the spatial location of the singing event, so the annotation was based only on the acoustic features of the song. We then applied the same annotation to all other audio clips from the same singing event. Note that due to this procedure, the annotator may not have been able to discern the individual identity of an Ovenbird in the more distant recordings from a singing event, but the annotation of the closest clip is still copied to the more distant clips. If the reviewed clip did not contain an Ovenbird song or the spectrogram did not contain sufficient detail to identify the song variant, the singing event was discarded and excluded from further analysis. We assessed annotator agreement rate by aligning the two reviewers’ clusters with majority voting (details in SI III).

### Training deep-learning feature extractors for individual identification

We train convolutional neural networks on spectrograms generated from the training dataset samples using a supervised classification approach, treating the survey location of the Ovenbird song clip as the class label. In this paradigm, the model must learn to correctly assign a song to the recording location where it was recorded. To use the trained model for acoustic identification of novel individuals, we remove the final fully-connected layer and use the output of the penultimate layer as the feature vector for clustering.

We use OpenSoundscape^46^ to train convolutional neural networks that take spectrograms as input. We use the default hyperparameters, preprocessing, and augmentation settings from OpenSoundscape version 0.12.0 CNN class, with the following modifications. We resample the audio to 32 kHz and normalize the audio signal. We crop spectrograms to 2-10 kHz, the frequency range of Ovenbird songs. We remove the random affine and frequency mask augmentations and add an augmentation that randomly shifts the start of the training sample by up to 0.5 seconds. We use overlay augmentation, which performs an averaging of the original sample with an additional sample not containing Ovenbird song, with a probability of 75% and weighting of the overlaid sample randomly selected between 0.01 and 0.6. Training scripts, experiment configuration files, and results are included in the GitHub repository.

We experimented with various loss functions for supervised classification models. We compared cross-entropy loss, binary cross-entropy loss, and sub-center ArcFace loss^47^ with 1, 2, 4, and 8 subcenters per class. ArcFace loss is a metric loss designed to maximize the angular separation of classes in the feature space. Sub-center arc-face loss generalizes this strategy by allowing a single class to cluster into multiple sub-centers in the feature space, which mirrors the structure of our training data, in which multiple individuals occur at a single survey location.

We also compared the supervised classification approach to a contrastive metric learning strategy. Contrastive metric learning losses are designed to teach the deep learning model to learn discriminative features between samples without providing the model with class labels, and may be well-suited for tasks like individual identification that require discriminating between novel categories. We designed a contrastive loss function that incentivises similarity or dissimilarity of pairs of samples in each mini-batch based on whether the samples were recorded at the same location (encourage similarity) or different locations (encourage dissimilarity). The loss function additionally incentivises the similarity of feature vectors for copies of the same sample that have been stochastically augmented two or more times.

Optionally, we include a self-supervised term of the loss function that periodically clusters the training set samples, then incentivizes the similarity of features for clips belonging to the same cluster. Details of the contrastive learning approach are given in SI VI.

We ran a series of model training experiments to compare training approaches and optimize hyperparameters. We ran univariate experiments, varying one parameter while holding all others constant at a base configuration, and training on a subset of 10,000 samples from the full training data for computational tractability. We ran experiments on the Bridges2 supercomputing resource^48^. The base configuration used supervised classification with binary cross-entropy loss and a ResNet18 architecture. Because model training and HDBSCAN clustering are stochastic, we ran each experimental configuration five times and report the mean and range of performance metrics across runs. We trained all models for 10 epochs with the AdamW optimizer^49^ with a mini-batch size of 128, learning rate of 0.005, and a cosine annealing learning rate schedule. Backbone experiments compared the ResNet18 and ResNet50 architectures with the HawkEars bird species classification model. We warm-start ResNet architectures by loading weights trained on the ImageNet image classification benchmark^50^ and using the pre-trained weights of HawkEars v0.1.0 when using the HawkEars backbone.

We also ran experiments to compare audio clip durations of 1, 2, 3, and 4 seconds; adding noise reduction (noisereduce Python package^51,52^) to audio preprocessing; and adding overlay augmentation in which an audio clip not containing Ovenbird is averaged with the clip containing Ovenbird during preprocessing. For the contrastive learning approach, we ran experiments with and without the self-supervision (SSL) strategy of intermittently clustering training samples to create pseudo-labels, and compared mini-batch sampling strategies that included 8 or 32 unique points per mini-batch (ppb).

We also investigated how model performance varied with the amount of training data. We trained models while varying the total dataset size (100, 500, 1000, 500, 10,000, 50,000, or all 94,378 samples) and while varying the number of unique survey points included in training data (8, 16, 32, 64, 128, or all 234 points).

We evaluated the performance of each experiment configuration on the validation set consisting of Ovenbird songs from 10 acoustic localization arrays. Based on these results, we then selected a configuration for training a full, final feature extractor. The full model used the supervised classification approach with 3-second audio clips as input, used a ResNet-18 backbone, trained with cross-entropy loss, included noise reduction and overlay in preprocessing, and was trained on the full training dataset. Performance saturated after one epoch, and we use a checkpoint from the end of the first training epoch for downstream tasks. The trained model is also included in the GitHub repository.

### Benchmarking against existing feature extraction models

We compare our trained feature extractors to state-of-the-art pre-trained models for bioacoustic feature extraction. Acoustic feature extraction using pre-trained deep learning methods (or other methods) may be sufficient for automated acoustic identification if the feature vectors extracted by the models capture the variation relevant to distinguishing individuals by song^20^. We first test the BirdNET^53^ and HawkEars^45^ bird species classification models. We also test the ResNet18 architecture pre-trained on the ImageNet image classification benchmark^50^. Similarly to above, we use the output of the penultimate layer as the feature embedding vector, which is the standard approach in representation learning for using CNNs for acoustic feature extraction^6,54^ and is the embedding approach used by BirdNet and HawkEars.

### Dimensionality reduction and clustering

Given embedding vectors for a set of Ovenbird song audio clips, we assign groups of clips to individual Ovenbird clusters using clustering algorithms. Before clustering, we reduce the dimensionality of feature vectors with manifold-based dimensionality reduction techniques (e.g., UMAP or t-SNE), which enables effective clustering of high-dimensional embeddings^31^. We then cluster all embedded Ovenbird detections using HDBSCAN^55^, a density-based hierarchical clustering algorithm that does not require specifying the number of clusters and includes a “noise” class of samples not allocated to any cluster. We standardize all feature dimensions (to mean 0, standard deviation 1) before dimensionality reduction and again before HDBSCAN clustering. The HDBSCAN “minimum number of samples” parameter affects the total number of clusters produced, with lower values resulting in more clusters. We set the HDBSCAN minimum number of samples parameter to 7 when clustering samples from the Localization Dataset and 30 when clustering samples from the Longitudinal Dataset because smaller values frequently produced unreasonable numbers of clusters per point (i.e., >20 clusters per sampling location in the Longitudinal Dataset, Figure S13). We leave the optimization of clustering parameters as a topic for future work.

We compare the performance of UMAP^30^ and t-SNE^29^ dimensionality reduction algorithms on the validation set. We test UMAP with 2,3,5,10,20,30 dimensions, and t-SNE with 2 and 3 dimensions, because the default t-SNE algorithm does not support creating more than 3 dimensions. Because these algorithms are stochastic, we repeat each experiment 30 times and report the mean and standard deviation of accuracy on the validation set.

### Evaluating automated individual identification performance

We evaluated automated individual identification approaches in two ways. First, we assessed how well automatically produced clusters match human-annotated clusters. Second, we evaluated how individual identification performance changed when an increasing number of individuals were simultaneously clustered.

The first evaluation assessed individual category discovery identification *per se*, that is, whether the model can accurately cluster a set of audio clips into one category per individual. We evaluated each feature extraction approach based on clustering performance on individual Ovenbirds from the 13 localization arrays (10 in the validation set, 3 in the test set). To increase the difficulty of the evaluation, we simultaneously clustered songs from all 10 validation grids or 3 test grids and did not provide any information about which grid the songs originated from. We evaluated the model only on the annotated Ovenbird singing events, rather than on all acoustic data from the localization arrays. Compared to a real-world application of the model, this evaluation did not include the effects of imperfect automated species recognition and focused instead on the individual classification problem.

We report accuracy and four clustering metrics that evaluate clustering performance versus ground truth: Adjusted Rand Index (ARI), Normalized Mutual Information (NMI), Feature Mutual Information (FMI, and V-measure (the harmonic mean of homogeneity and completeness)^56^. Computing the overall song-level accuracy requires first creating a mapping between the proposed clusters created by the algorithm and the annotated clusters labeled during manual review, where the algorithm may produce more or fewer clusters than the annotations have. Similarly to Zhao et al.^15^, we use the Hungarian algorithm (SciPy^57^ package’s optimize.linear_sum_assignment) to find the one-to-one mapping of proposed clusters to annotation clusters that results in the highest song-level accuracy (fraction of correctly classified songs). Songs in proposed clusters that are not mapped to any of the annotation clusters are considered to be incorrectly classified, as are songs from annotated clusters that do not belong to any proposed cluster.

We also report three measurements of recall. Automated species recognizer recall is the fraction of Ovenbird songs that are retained after thresholding and post-processing the results of the automated species recognition model, and is evaluated using the annotated set of 1,000 randomly selected clips from the Localization Dataset. Individual identification recall is the fraction of detected songs assigned to a cluster (some songs are left unclustered when using the HDBSCAN clustering algorithm). End-to-end recall is the product of these and equals the fraction of all songs in the passive acoustic monitoring data that are assigned to an individual.

Second, we evaluate automated individual identification on a pool of an increasing number of individuals and sampling locations. In other words, how does performance when separating a handful of individuals from a single location compare to performance when separating hundreds of individuals from hundreds of locations? Typically, individual clustering performance will decrease as the total number of true clusters increases.

For this analysis, we do not use human-annotated labels. Instead, we select Ovenbird song clips from a passive acoustic monitoring study (see below), and filter to a set of ARU sampling locations with a minimum of 1 km spacing (n=252) to minimize the likelihood of an individual being detected at multiple locations. We first randomly select N (2, 4, 8, 16, 32, 64, or 128) points and retrieve the dimensionality-reduced Ovenbird song embeddings from the trained feature extractor. If there are more than 200*N songs, we randomly subset to 200*N songs for computational tractability. We perform unsupervised clustering of the embeddings and evaluate two metrics. First, we measure cluster contamination, which we define as the fraction of songs included in a cluster that are from a sampling location different from the most-represented sampling location in the cluster. For example, if a cluster has 8 samples from Point A, 1 from Point B, and 1 from Point C, the cluster contamination is 0.2. Under the assumption that the same bird does not occur at two points a kilometer apart, cluster contamination indicates that the model has incorrectly included songs from different individuals in a single cluster. Second, we compare the number of clusters discovered when all songs are simultaneously clustered over the number of clusters discovered when songs from each sampling location are separately clustered, an assessment of recall at the level of individual birds.

### Study site

State Game Lands 34 and Ohiopyle State Park are distributed across the North-central and Southwest regions of Pennsylvania, USA, respectively. Forests within both of these sites range from 15-130 years old and consist of mixed-oak and northern hardwood forest types (see Lyon et al.^40^ for more information).

Sampling points were generated within the two study sites using the “create random points tool” in ArcGIS 10.4, with points stratified by forest age class based on inventory data collected by either the Pennsylvania Game Commission or the Pennsylvania Department of Conservation and Natural Resources (See Fiss^58^ for additional details). We separated survey points by a minimum of 250m to avoid double-counting individuals following suggestions from Ralph et al.^59^ and required that points be situated greater than 100m from any roads or trails.

Microhabitat data were collected in situ at each point in either 2020 or 2021. Microhabitat variables included basal area, sapling and shrub stem density, and leaf litter percent cover (see Fiss 2023 for information on how each variable was collected). Hill aspect and slope were calculated for each sampling point using the FedData^60^ package in R. Additionally, we extracted percent canopy cover for each point from a canopy cover data source generated in 2021 (NLCD https://www.mrlc.gov/viewer/). Using QGIS^61^, we calculated the average canopy cover value of all 30 m pixels within a 50m radius of each point.

The localization arrays were located within the Central Mixedwood Natural Subregion of the Boreal Forest Natural Region of Alberta^44,62^. Each localization array was set up surrounding a reclaimed wellsite (typically 1 ha area of shorter woody vegetation than surrounding forest) and adjacent linear feature, with no significant additional disturbance (i.e., forestry harvest) in the area sampled. The wellsites were within mature (∼80 years old) trembling aspen (*Populus tremuloides*) and balsam poplar (*Populus balsamifera*) dominated forests, and were selected to sample a range of woody vegetation regeneration following reclamation (Wilson and Bayne 2018).

### Ovenbird population density and annual survival in Pennsylvania

We conducted a non-invasive individual resighting study of Ovenbirds in the four-year PAM dataset from Pennsylvania by using our individual identification approach to discover and resight individuals based on their acoustic signature. Using this resighting data, we analyzed Ovenbird abundance and annual survival across the Longitudinal Dataset. For these analyses, we subset the acoustic monitoring dataset to a 12-day window from May 15-26 each year, which corresponds to the Ovenbird breeding period in these study regions^34^. Thus, we focus on the breeding period in which Ovenbirds are likely to be defending a stable territory, and are likely more stationary than during migration, territory settlement, and post-breeding periods. We analyzed a 1.5-hour window from 6:00 AM to 7:30 AM on each day. For survival and abundance analyses, we only included the 126 survey locations with complete coverage of these dates and times across all 4 years (2021-2024).

Ovenbirds have very high site fidelity across their lifetime, typically establishing territories that overlap completely or partially with their previous year’s territory^25^. In a focused search across neighboring pairs of survey points (126 points, spacing 258-1695 m, median 581 m) and four years of data, we found that only 3 of 405 individual Ovenbirds also occurred in more than one audio clip at the closest neighboring point (details in SI V). An additional 6 individuals occurred in exactly one 3-second audio clip from the nearest neighboring point. Therefore, we treat points as independent in the survival analysis and do not consider the possibility of an individual occurring at multiple points.

### Generating automated detections of individuals

To generate automated individual Ovenbird detections across this dataset, we use the HawkEars species classifier and the post-processing steps described above. When aggregating over increasing numbers of 3-second clips, the precision of species classification will decrease, and recall will increase. Therefore, since PAM data generates many clips for every sampling occasion, we favor precision over recall in the case study by using a higher species classification threshold score for Ovenbird (logit score 1.0, precision 0.97, recall 0.69). We use a threshold of 0.0 for rejecting clips where other classes were detected by HawkEars.

Because we assume individuals do not occur at multiple points, we perform automated clustering of Ovenbird song embeddings on a point-by-point basis. Compared to clustering embeddings from all songs from the entire dataset simultaneously, clustering one point at a time resulted in finding more distinct individuals (Figure 5d). We observed that dimensionality reduction (which precedes clustering) was more effective when performed across the entire dataset versus on a point-by-point basis. Therefore, we performed dimensionality reduction on the entire dataset, then clustering on detections from one point (and all four years) at a time.

This process results in a set of clusters for each point, with each cluster containing a set of audio clips that the model believes share similar individual-level Ovenbird song features.

### Reviewing automated detections to create a detection history

We reviewed the clusters generated with the previous steps to create a human-verified detection history of individual recaptures using a two-phase review process. In Phase 1, we removed heterogeneous clusters and merged clusters representing the same individual by inspecting audio clips from automatically generated clusters on a per-point basis. These clusters contained songs from all 4 study years. We randomly selected ten audio clips from each cluster and inspected their spectrograms visually. If the audio clips did not contain an Ovenbird song or contained distant songs in which the characteristics of individual songs could not be determined, the cluster was removed from further analysis. If a single, visually consistent Ovenbird song type occurred in at least 5 of the 10 audio clips and was distinct from other song types in previously reviewed clusters, we marked the clips containing that song type as Representative Songs for the cluster and gave the cluster a unique label (e.g., RK001_1 for a cluster detected at the point named RK001). Based on the results of the evaluation on the Localization Dataset, we consider each of these clusters to represent a single individual male Ovenbird’s song. If the song type occurring at least five times matched a previously established cluster, we merged the clusters. If no song type occurred in at least 5 of the 10 clips, we considered the cluster to be heterogeneous and removed it from further analysis.

In Phase 2, we created a detection history for each of 12 days (15-26 May) on each year (2021-2024) for each point and each individual by reviewing detections from the clusters resulting from the Phase 1 review. For each cluster, day, and year, we selected the single audio clip that had the highest feature similarity to any of the clips reviewed and included in the cluster during Phase 1. We visually inspected these samples to confirm whether the individual represented by the cluster was present, resulting in a presence-absence detection history for each day, year, and cluster.

### Cormack-Jolly-Seber model of apparent survival

We statistically estimate annual apparent survival of male Ovenbirds from our individual resighting histories by fitting a hierarchical Cormack-Jolly-Seber (CJS) model with spatial replication^63^. The CJS model estimates two parameters, annual apparent survival (phi) and annual detection probability (p), based on recapture histories of individuals from repeated sampling occasions. In this model, permanent emigration cannot be distinguished from death, and apparent survival is thus interpreted as the probability of both surviving and returning to the site one sampling period (e.g., year) later. Apparent survival thus provides a lower bound on true annual survival. The annual detection probability incorporates both imperfect detection (the bird used the point but was not detected) and temporary emigration (i.e., the bird did not use the point during one year). Here, we call the survey locations of each ARU point and the two geographically separated landscapes (SGL 34 and Ohiopyle) blocks to avoid the overloaded term site.

In its basic form, the CJS model assumes that after first being detected (and alive, z_0=1), individuals survive from one survey period to the next with some apparent survival probability phi (and once dead, remain dead):

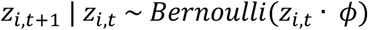

Where z is 1 if individual is alive at time t and 0 if dead; *i* indexes individuals; *t* indexes sequential survey periods; and ф is the apparent survival. The model assumes that no misidentifications or false-positive identifications of individuals occur. We fit models with constant survival and with survival varying by year. On each survey period, alive individuals are detected with probability *p*, while dead individuals are not detected:

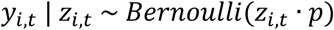

Where *y_i,t_* is 1 if individual i is observed on survey occasion *t* and 0 if it is not observed.

Due to a very low rate of cross-point recaptures (see above and SI V), our data were not suitable for a spatial capture recapture model, in which recaptures of an individual across spatial sampling points inform the estimation of a movement or dispersal kernel. Instead, in our CJS model, we treat each survey location (N=126) as independent and assume that individuals only occur at a single point.

We aggregate our 12-day detection histories into yearly detection histories for each individual. Although the CJS has been adapted for robust-design surveys such as our 12-day, 4-year detection history, we leave the application of robust-design survival models to passive acoustic resighting data as a topic for future work. Because the occurrence of transient individual birds who occur only briefly at a sampling location before permanently emigrating can result in lower estimates of apparent survival^63^ (chapter 15), we fit survival models with and without transient individuals. We adopted an ad hoc approach to separate transient and resident individual male Ovenbirds in our study, by considering birds detected on at least 4 days of a 12-day detection history within a year to be residents.

In addition to fitting the basic CJS model, we investigate differences in apparent survival across habitat types and management regimes by including point-level covariates on apparent survival. We use JAGS via the jagsUI package^64^ in R to define a Bayesian formulation of the model in R and run Markov Chain Monte Carlo (MCMC) sampling, following the example provided in Kéry and Royle^63^ (section 3.4.3). Since individuals are nested within points, and points within regions (Ohiopyle and State Game Lands 34), our models include a random effect on point and a fixed effect for region. We also expect that survival might vary by year due to unobserved conditions such as weather during migration or habitat conditions on the breeding grounds. Therefore, our models allow apparent survival phi to vary by year. Thus, apparent survival phi in the null model is indexed by point (s) and year (t):

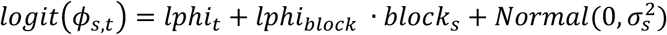

Where *block*_s_ equals 0 for points at Ohiopyle and 1 for points at SGL 34. We assume detection probability p is constant across points, blocks, and years, so the detection process of the model remains the same as above.

Based on a literature review of Ovenbird habitat selection^34^, we selected percent canopy cover, basal area (a measure of tree size and density), and ground slope as variables that might explain variation in apparent survival across points. Our full model includes these variables as additive predictors for apparent survival phi:

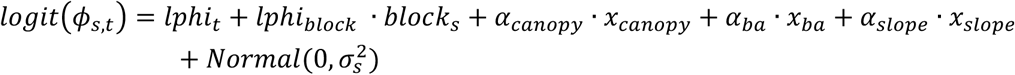

Where *X_canopy_* is the standardized percent canopy cover, *X_ba_* is the standardized basal area, and *X_slope_* is the standardized ground slope.

### Generalized Linear Models of Abundance

We fit generalized linear models to investigate whether Ovenbird abundance was explained by habitat covariates and how abundance varied over time. We directly model the number of observed individuals as the response variable, leaving hierarchical models of abundance with stochastic detection processes as a topic for future work. We model abundance (number of individuals per point) as a Poisson process with rate lambda. We model lambda with the same point-level covariates as the full CJS model (percent canopy cover, basal area, and slope), and intercept effects for year and block (SGL 34 versus Ohiopyle), with a log link function:

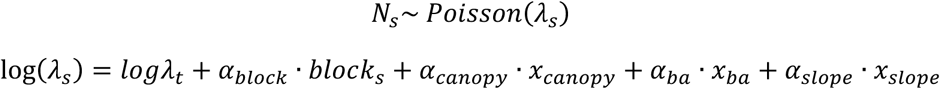

Where the intercept logλ*_t_* indexes over years, and *block*_s_ is 1 for points at SGL 34 or 0 for points at Ohiopyle. We also fit all sub-models with and without each covariate and compare AICc using the R package MuMIn^65^.

