## Supplementary Materials for "Automated identification of individual birds by song enables multi-year recapture from passive acoustic monitoring data"

### Supplementary Information

#### Supplementary Figures

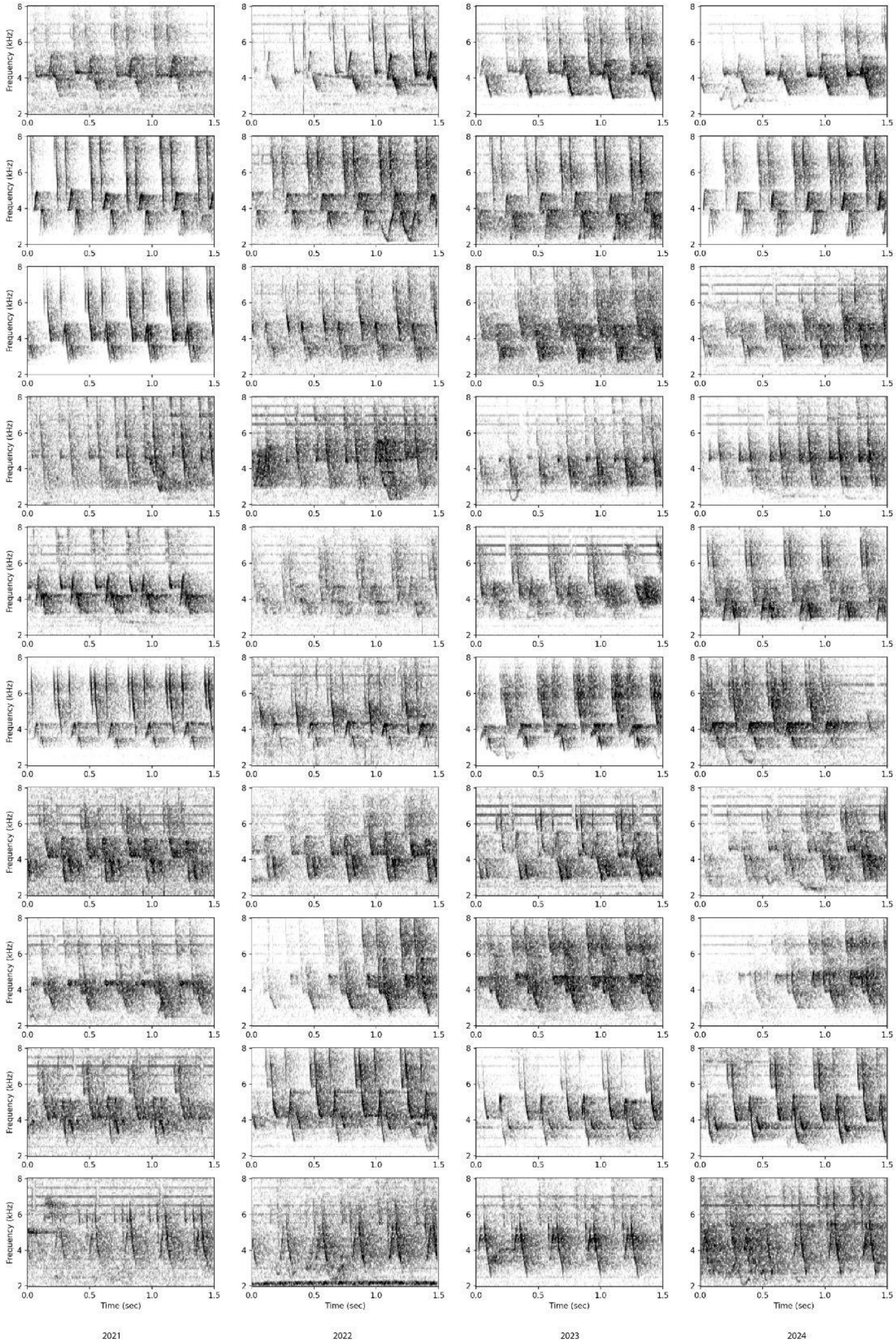
Frequency (kHz)
Time (sec)
2021
2022
2023
2024

Figure S1: Ovenbird songs contain stable acoustic signatures that persist across years without apparent changes. Each row shows spectrograms from each year 2021-2024 for a randomly selected individual from the Longitudinal Dataset. Log-valued, linear-frequency spectrograms were created from normalized 1.5-second audio clips using a 512-sample Hanning window with a step size of 256 samples and show from -55 (white) to -10 (black) dBFS using OpenSoundscape.

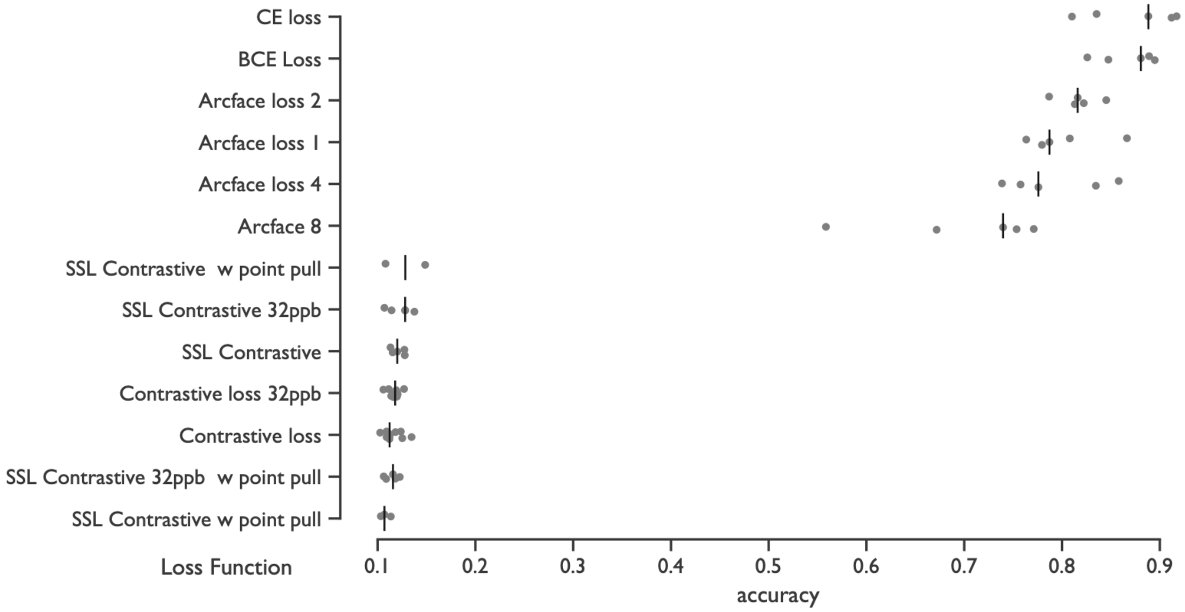


Figure S2: Validation set accuracy of each loss function from five training experiment runs. Points show the individual run values, lines show the median. The supervised classification training approach with cross-entropy (CE) or binary cross-entropy (BCE) loss performed best. Numbers after Arcface loss indicate the number of subcenters. Contrastive loss was trained with and without “point pull”, where samples from the same point were pulled together in the feature space, and with 8 points per batch (ppb) unless noted. Self-supervised learning (SSL) approaches use feature clustering to create pseudo-labels on the training set (see Methods).

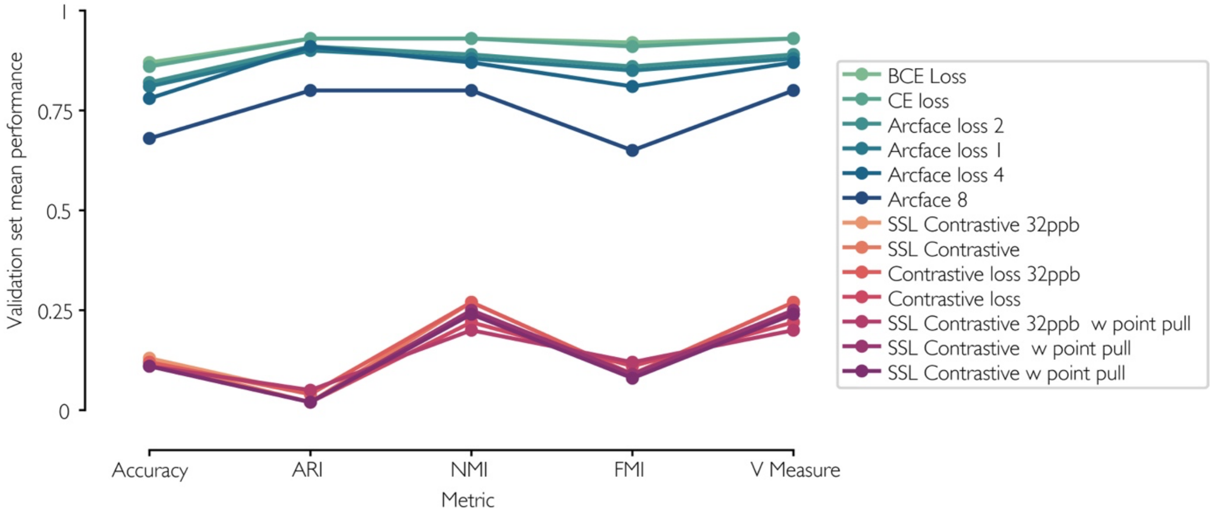


Figure S3: Comparison of model performance for different loss functions. Points show mean validation set performance of loss functions for each metric. See Methods for definitions of loss functions.

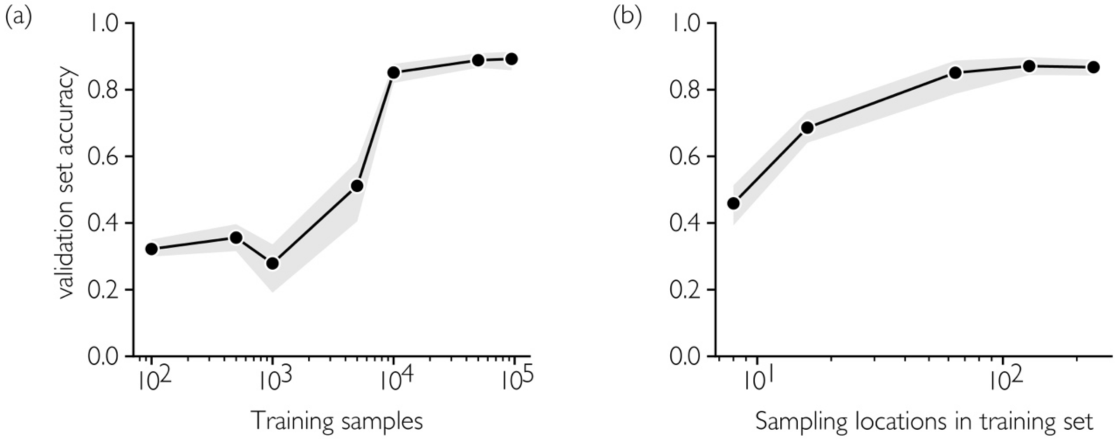


Figure S4: Validation set accuracy increased with a saturating relationship against (a) the number of training samples and the (b) the number of unique sampling locations in the training data. The shaded regions show a 95% confidence interval on the mean across 5 runs.

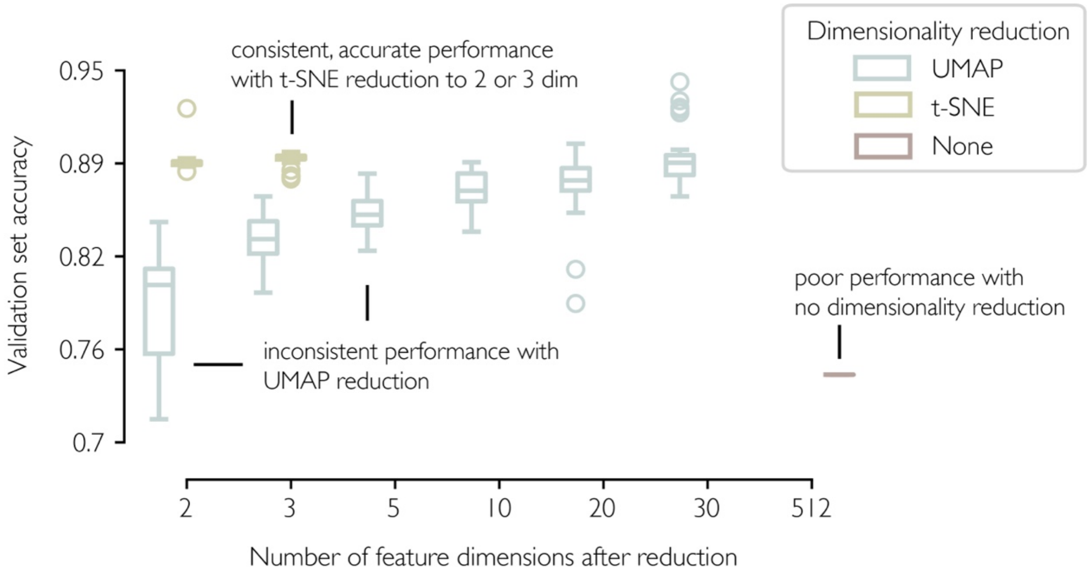


Figure S5: Validation set accuracy for 30 runs of each dimensionality reduction configuration

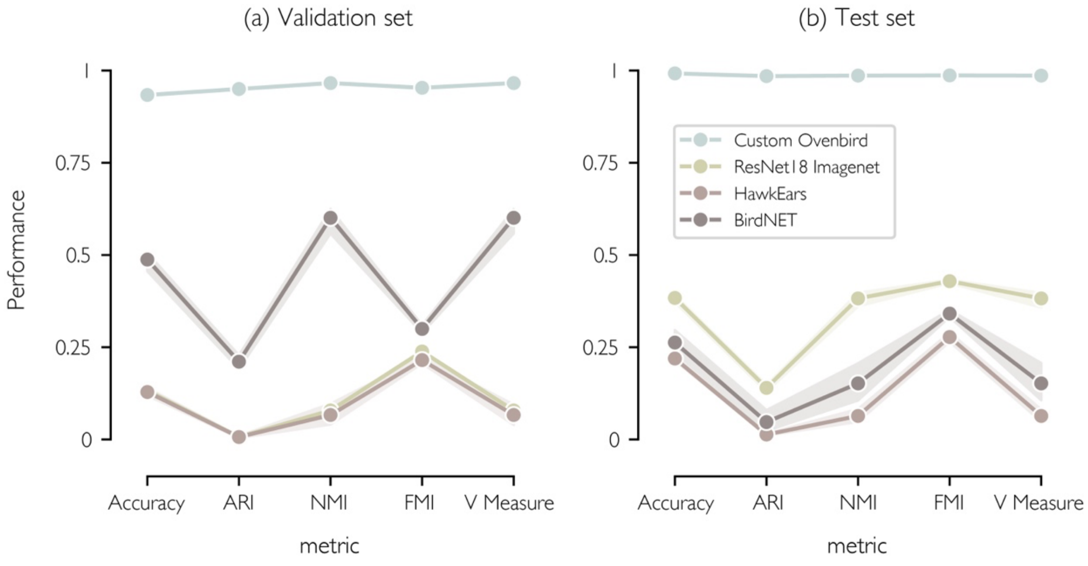


Figure S6: Validation and test-set performance of our feature extractor trained to discriminate individual Ovenbirds compared to features from pretrained models (shaded area shows 95% confidence interval on the mean based on 30 runs of stochastic clustering).

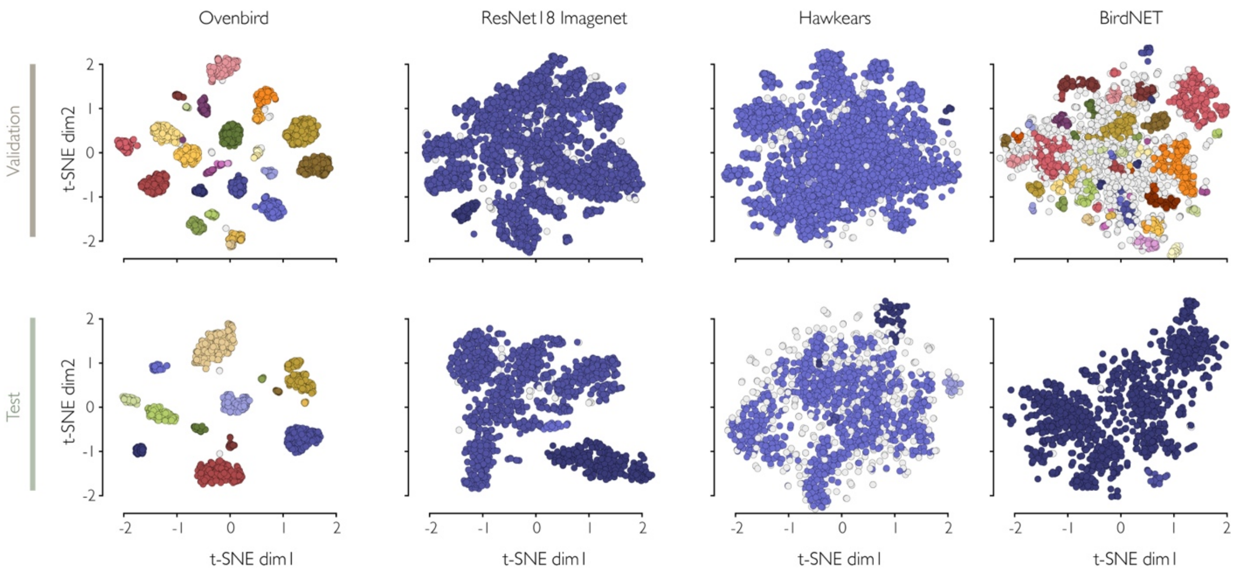


Figure S7: Validation and test set embeddings colored by the automated cluster assigned with HDBSCAN. Grey points are songs not assigned to any cluster. Feature vectors were generated using each of four feature extractors (columns) and reduced to 2 dimensions with t-SNE.

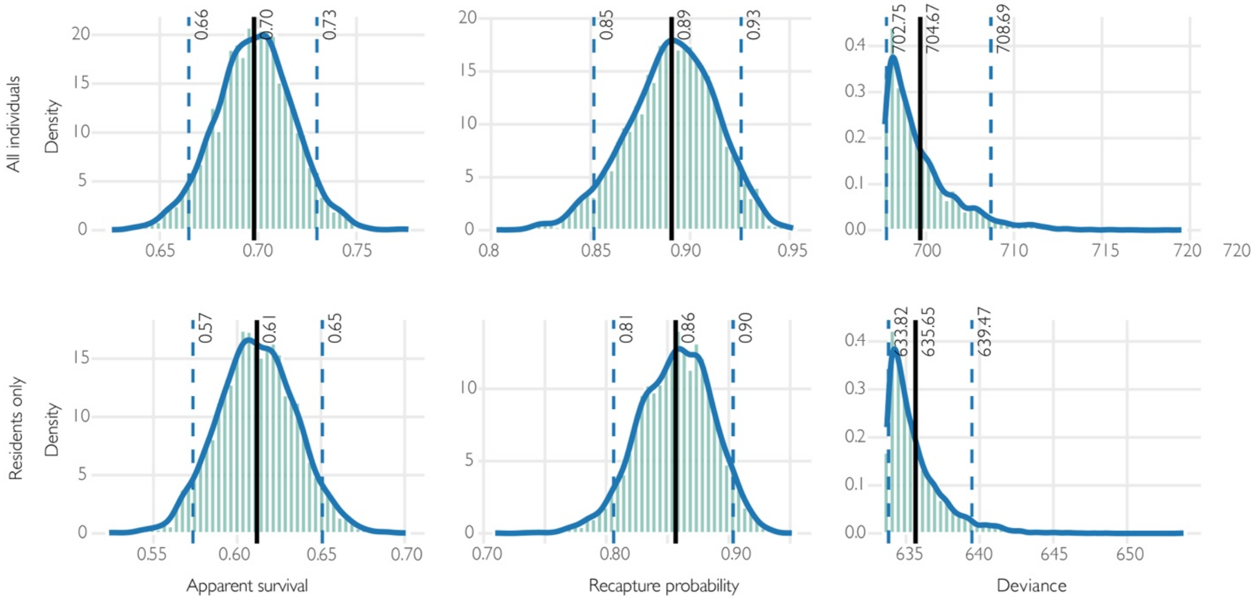


Figure S8: Base CJS parameter posterior distributions for models with all individuals and only residents

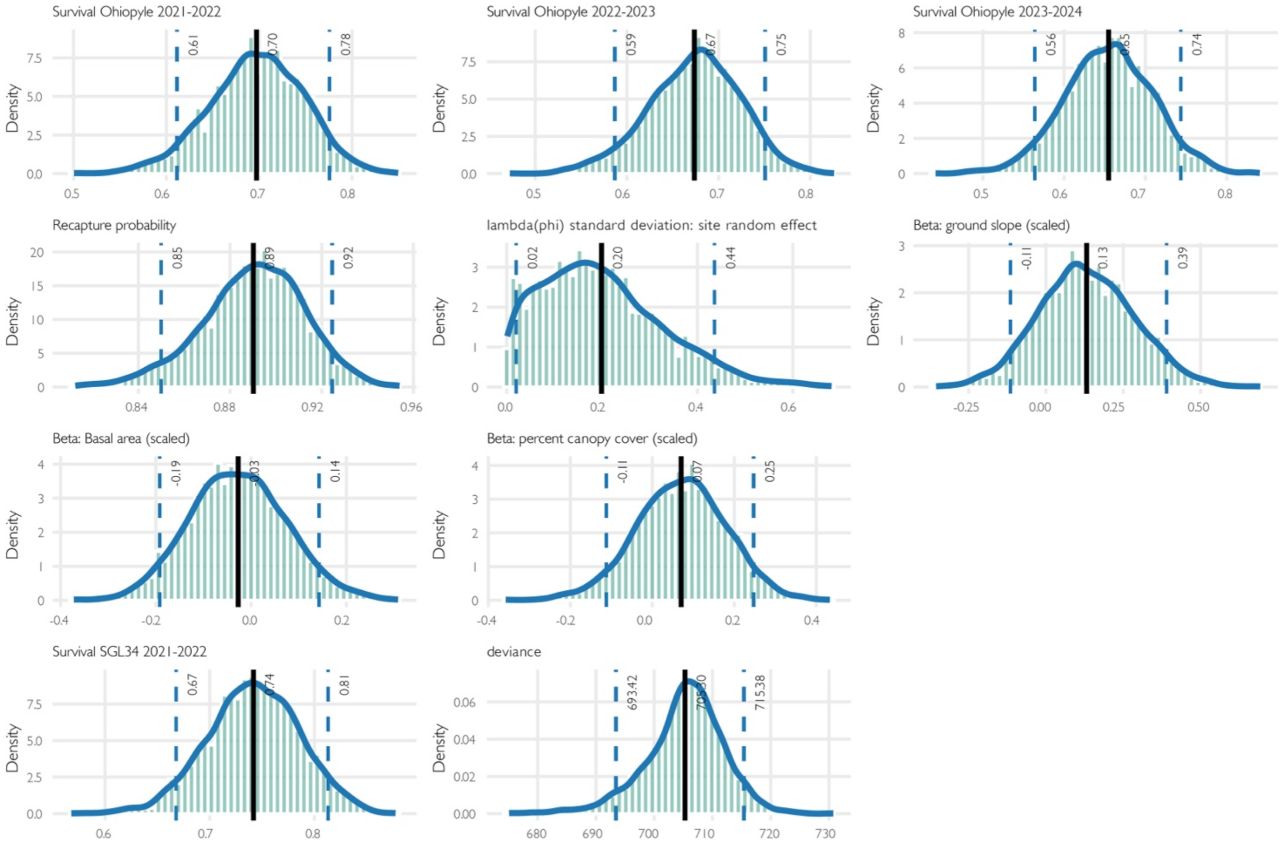


Figure S9: posterior distributions of parameters with the full CJS model with all individuals. Vertical lines show the 5%, mean, and 95% values of the posterior.

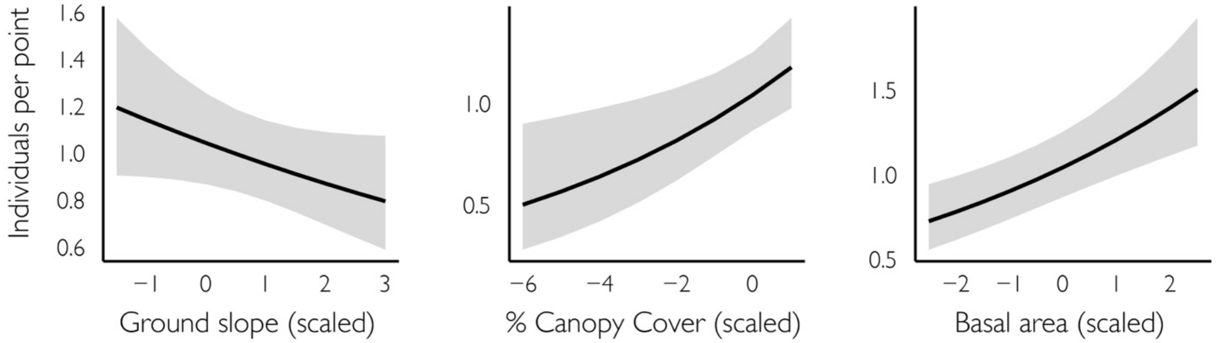


Figure S10: Partial effect of each habitat covariate on Ovenbird abundance in units of standard deviations from the mean. Shaded regions show 95% confidence intervals on the modeled mean.

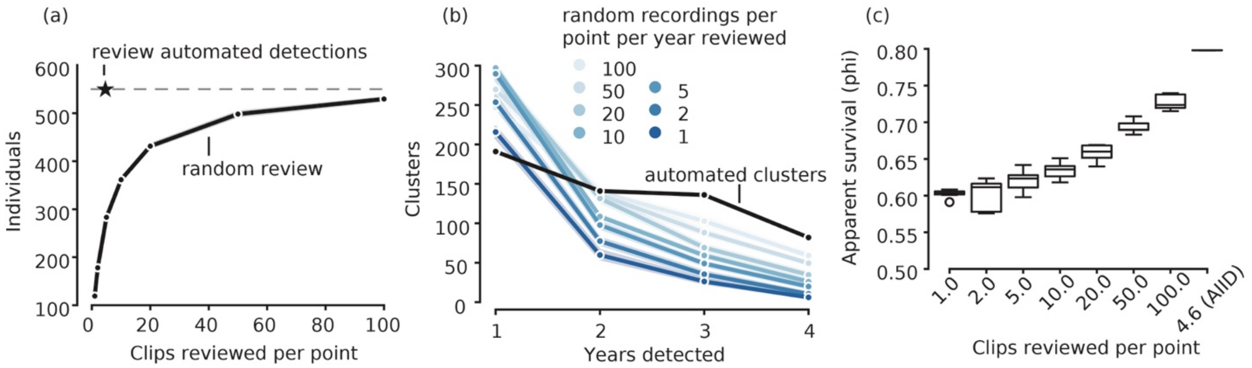


Figure S11: Simulations comparing abundance and survival estimation using automated individual identification versus annotation of randomly selected songs. (a) Automated individual recognition greatly increases the efficiency of finding individuals (star) compared to reviewing randomly selected clips from each survey point (circles); (b) individuals are resighted on more years with automated individual recognition than with random review; (c) as a result, estimates of apparent annual survival are biased low and are highly variable under the random sampling strategy, compared to estimates using automated individual recognition.

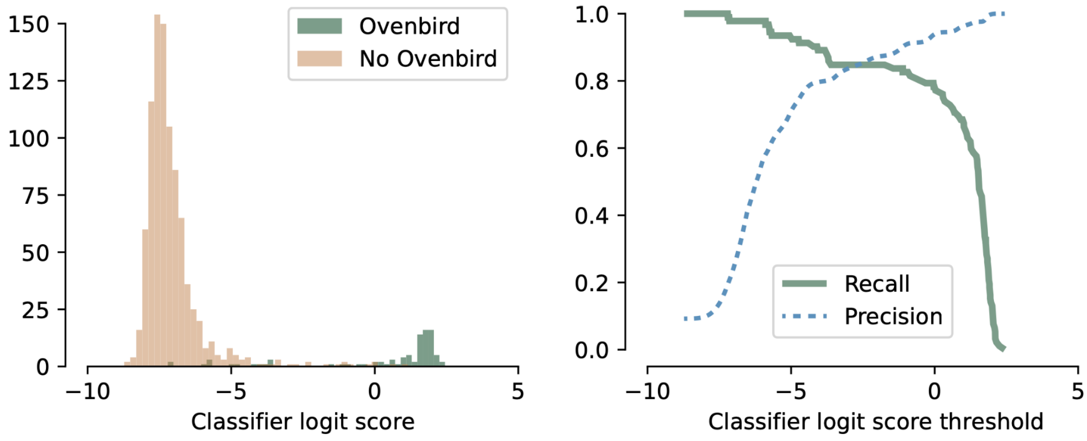


Figure S12: HawkEars performance for Ovenbird song detection on an annotated set of 1000 randomly selected clips from the Localization Dataset. (a) Histogram of classifier logit scores for 3-second clips with and without Ovenbird; (b) precision and recall versus score threshold

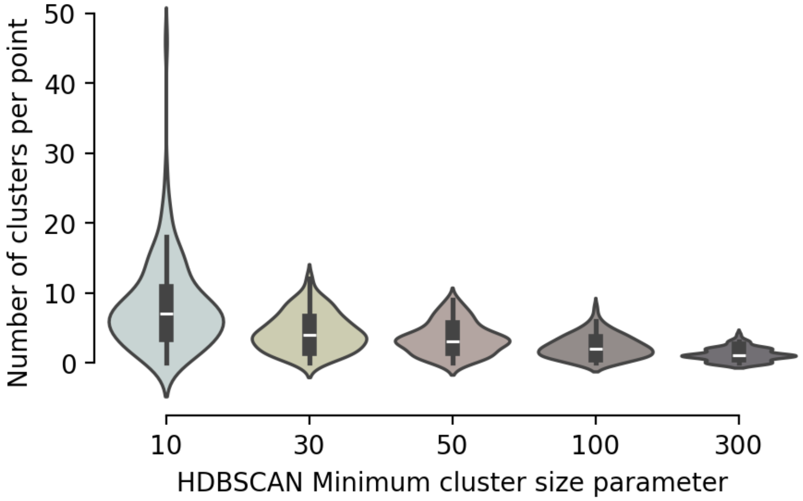


Figure S13: Violin plot of the number of clusters generated per point in the Longitudinal Dataset, for different values of HDBSCAN minimum cluster size parameter

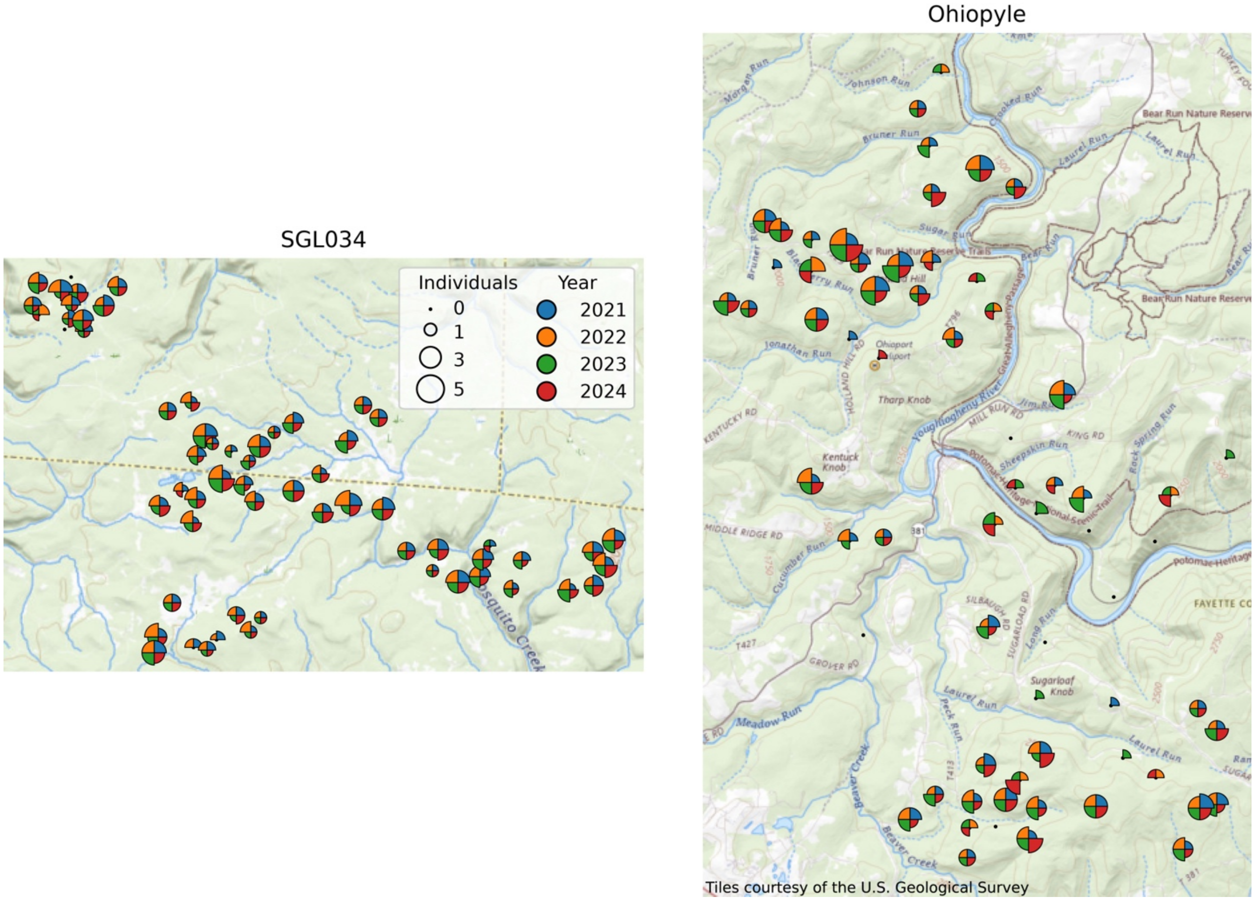


Figure S14: Maps of Ovenbird individuals detected per year at each point, with size of wedges scaling with number of individuals detected. Black dots indicate points where no Ovenbird songs were detected that year.

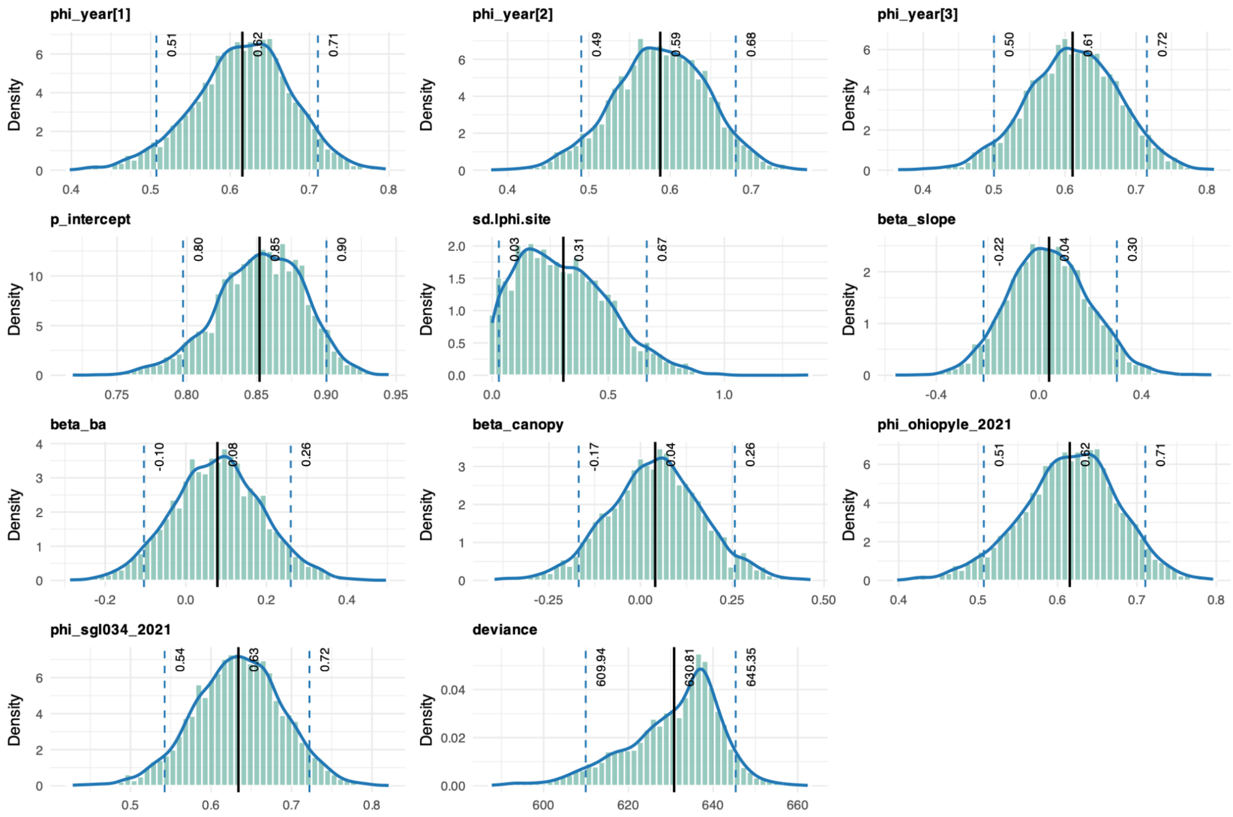


Figure S15: Posterior distributions of parameters with the full CJS model with only resident individuals. Vertical lines show the 5%, mean, and 95% values of the posterior.

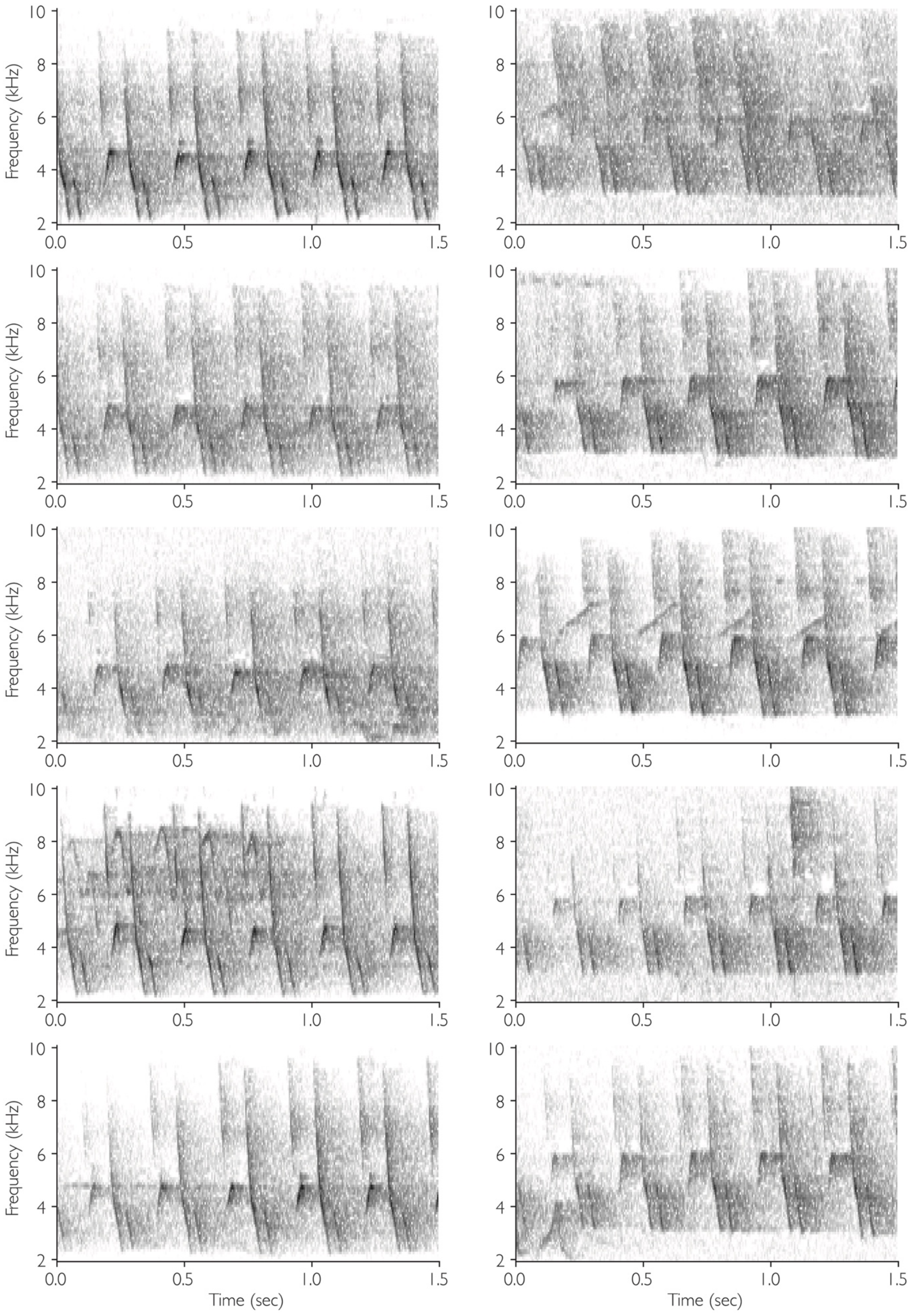
Frequency (kHz)
Time (sec)
10
8
6
4
2
0.0
0.5
1.0
1.5

Figure S16: Song examples from two individuals (one individual per column). One annotator initially marked songs from both individuals as being the same song variant, while the other annotated two separate individuals. Note that consistent difference in overall frequency, with the first individual having lower frequency content than the second.

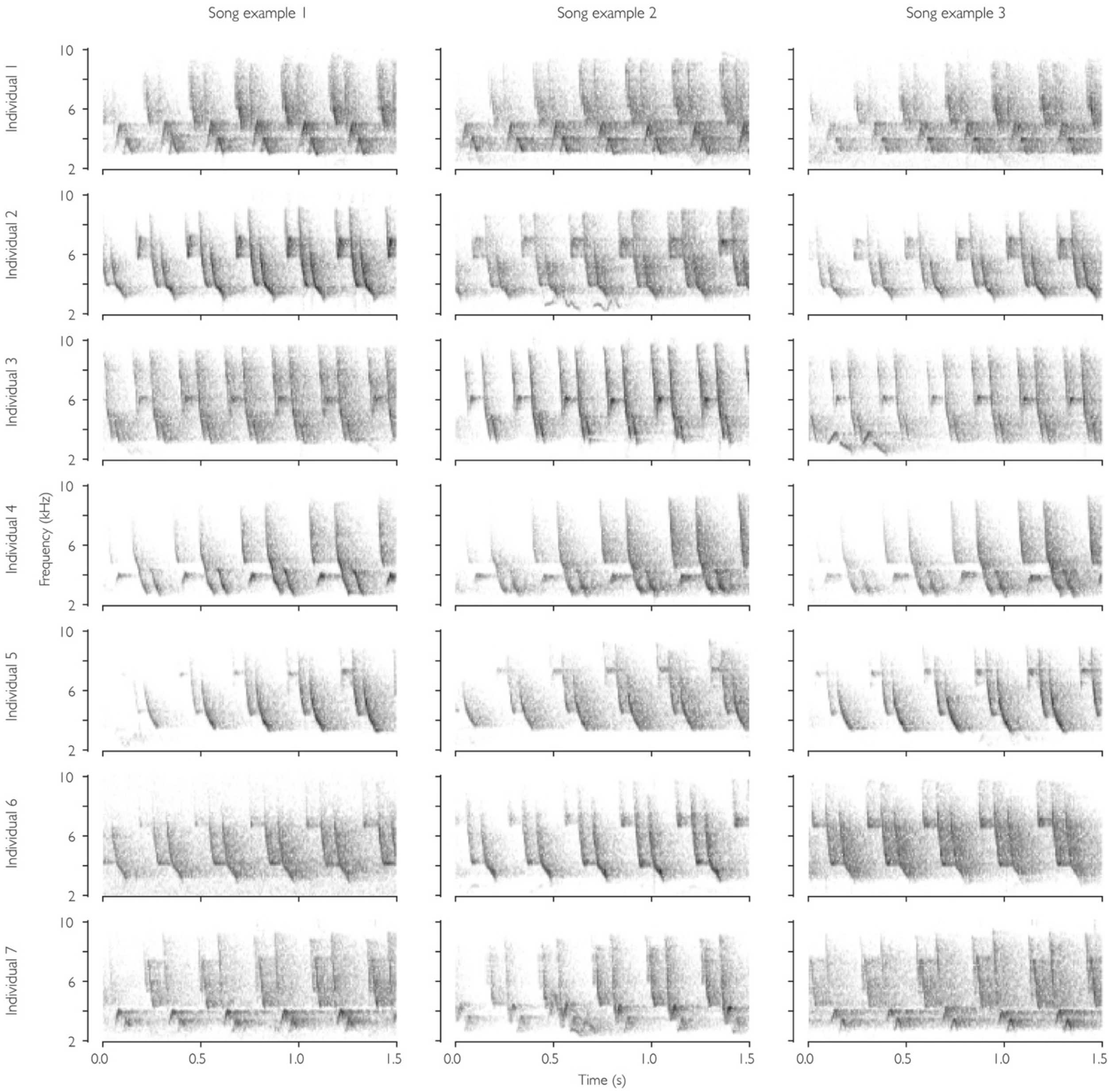


Figure 17: Song examples from two individuals (one individual per column). One annotator initially marked songs from both individuals as being the same song variant, while the other annotated two separate individuals. Note the subtle difference in the upward-sweeping note: on the left, it is centered at 4 kHz and disconnected from the previous note, while on the right, it is centered at 4.6 kHz and connected to the previous note.

#### Supplementary Tables

Table S1: Validation set performance for each training strategy and loss function.

Mean and range across 5 training runs are given for each metric (see body for definitions). The supervised classification training approach with cross-entropy (CE) or binary cross-entropy (BCE) loss performed best. Numbers after Arcface loss indicate the number of subcenters. Contrastive loss was trained with and without “point pull”, where samples from the same point were pulled together in the feature space, and with 8 points per batch (ppb) unless noted. Self-supervised learning (SSL) approaches use feature clustering to create pseudo-labels on the training set (see Methods).

| Loss approach | Accuracy | ARI | NMI | FMI | V-Measure |
|---|---|---|---|---|---|
| CE loss | 0.87 (0.81-0.92) | 0.93 (0.92-0.94) | 0.93 (0.91-0.96) | 0.91 (0.89-0.94) | 0.93 (0.91-0.96) |
| BCE Loss | 0.87 (0.83-0.89) | 0.93 (0.91-0.94) | 0.93 (0.91-0.95) | 0.91 (0.87-0.94) | 0.93 (0.91-0.95) |
| Arcface loss 2 | 0.82 (0.79-0.85) | 0.91 (0.88-0.93) | 0.89 (0.88-0.90) | 0.85 (0.84-0.88) | 0.89 (0.88-0.90) |
| Arcface loss 1 | 0.80 (0.76-0.87) | 0.90 (0.85-0.92) | 0.88 (0.85-0.90) | 0.85 (0.79-0.90) | 0.88 (0.85-0.90) |
| Arcface loss 4 | 0.79 (0.74-0.86) | 0.91 (0.91-0.92) | 0.88 (0.85-0.90) | 0.83 (0.76-0.89) | 0.88 (0.85-0.90) |
| Arcface 8 | 0.70 (0.56-0.77) | 0.80 (0.74-0.87) | 0.80 (0.73-0.84) | 0.67 (0.45-0.79) | 0.80 (0.73-0.84) |
| SSL Contrastive w point pull | 0.13 (0.11-0.15) | 0.03 (0.02-0.04) | 0.25 (0.24-0.25) | 0.10 (0.09-0.11) | 0.25 (0.24-0.25) |
| SSL Contrastive 32ppb | 0.12 (0.11-0.14) | 0.04 (0.02-0.06) | 0.22 (0.01-0.28) | 0.12 (0.09-0.24) | 0.22 (0.01-0.28) |
| SSL Contrastive | 0.12 (0.11-0.13) | 0.02 (0.01-0.02) | 0.26 (0.25-0.27) | 0.09 (0.09-0.10) | 0.26 (0.25-0.27) |
| Contrastive loss 32ppb | 0.12 (0.11-0.13) | 0.04 (0.01-0.08) | 0.24 (0.01-0.29) | 0.11 (0.09-0.23) | 0.24 (0.01-0.29) |
| Contrastive loss | 0.12 (0.10-0.13) | 0.02 (0.02-0.03) | 0.23 (0.02-0.27) | 0.10 (0.08-0.24) | 0.23 (0.02-0.27) |
| SSL Contrastive 32ppb w point pull | 0.11 (0.11-0.12) | 0.04 (0.02-0.07) | 0.21 (0.01-0.26) | 0.12 (0.09-0.23) | 0.21 (0.01-0.26) |
| SSL Contrastive w point pull | 0.11 (0.10-0.11) | 0.02 (0.02-0.02) | 0.24 (0.22-0.25) | 0.08 (0.08-0.08) | 0.24 (0.22-0.25) |

Table S2: Automated individual recognizer performance increases with the number of training samples

Average and range of automated individual vocal recognition performance on the validation set, across 5 repeated training runs. Each row shows the results of an experiment where the total number of samples included in the training set was restricted to a random subset of all samples. See text for definitions of the metrics reported in each column.

| Number of training samples | Accuracy | ARI | NMI | FMI | V-Measure |
|---:|---:|---:|---:|---:|---:|
| 100 | 0.32 (0.29-0.38) | 0.22 (0.18-0.25) | 0.48 (0.44-0.51) | 0.17 (0.15-0.19) | 0.48 (0.44-0.51) |
| 500 | 0.36 (0.28-0.41) | 0.14 (0.09-0.20) | 0.52 (0.43-0.56) | 0.18 (0.15-0.21) | 0.52 (0.43-0.56) |
| 1000 | 0.28 (0.11-0.35) | 0.14 (0.10-0.18) | 0.42 (0.05-0.54) | 0.18 (0.16-0.24) | 0.42 (0.05-0.54) |
| 5000 | 0.51 (0.31-0.60) | 0.42 (0.20-0.55) | 0.68 (0.52-0.77) | 0.44 (0.18-0.58) | 0.68 (0.52-0.77) |
| 10000 | 0.85 (0.76-0.89) | 0.93 (0.91-0.94) | 0.92 (0.88-0.95) | 0.90 (0.82-0.94) | 0.92 (0.88-0.95) |
| 50000 | 0.89 (0.84-0.92) | 0.94 (0.93-0.95) | 0.94 (0.92-0.96) | 0.93 (0.90-0.95) | 0.94 (0.92-0.96) |
| 94378 | 0.89 (0.83-0.92) | 0.95 (0.93-0.96) | 0.94 (0.91-0.95) | 0.92 (0.87-0.94) | 0.94 (0.91-0.95) |

Table S3: Automated individual recognizer performance increases with the number of unique sampling locations in the training data

Average and range of automated individual vocal recognition performance on the validation set, across 5 repeated training runs. Each row shows the results of an experiment where the total number of unique sampling locations included in the training set was restricted to a random subset of all sampling locations. See text for definitions of the metrics reported in each column.

| Number of unique sampling locations | Accuracy | ARI | NMI | FMI | V-Measure |
|---:|---:|---:|---:|---:|---:|
| 8 | 0.46 (0.35-0.51) | 0.51 (0.43-0.56) | 0.66 (0.61-0.71) | 0.43 (0.34-0.54) | 0.66 (0.61-0.71) |
| 16 | 0.69 (0.62-0.77) | 0.76 (0.72-0.78) | 0.82 (0.79-0.86) | 0.69 (0.63-0.77) | 0.82 (0.79-0.86) |
| 64 | 0.85 (0.73-0.89) | 0.92 (0.90-0.93) | 0.91 (0.86-0.93) | 0.87 (0.71-0.92) | 0.91 (0.86-0.93) |
| 128 | 0.87 (0.83-0.91) | 0.93 (0.92-0.94) | 0.93 (0.91-0.94) | 0.91 (0.88-0.93) | 0.93 (0.91-0.94) |
| 234 | 0.87 (0.83-0.89) | 0.93 (0.91-0.94) | 0.93 (0.91-0.95) | 0.91 (0.87-0.94) | 0.93 (0.91-0.95) |

Table S4: Comparison of deep learning backbones

Mean and range of automated recognition performance across 5 training runs  are given for each metric for three model architectures (see Methods).

| **Backbone** | **Accuracy** | **ARI** | **NMI** | **FMI** | **V-Measure** |
|---|---|---|---|---|---|
| ResNet18 | 0.87 (0.83-0.89) | 0.93 (0.91-0.94) | 0.93 (0.91-0.95) | 0.91 (0.87-0.94) | 0.93 (0.91-0.95) |
| ResNet50 | 0.80 (0.73-0.84) | 0.85 (0.69-0.92) | 0.90 (0.87-0.91) | 0.84 (0.72-0.89) | 0.90 (0.87-0.91) |
| HawkEars | 0.58 (0.48-0.66) | 0.51 (0.44-0.69) | 0.71 (0.64-0.78) | 0.47 (0.33-0.64) | 0.71 (0.64-0.78) |

Table S5: Mean validation set performance for training experiments comparing clip duration

Mean and range of automated recognition performance across 5 training runs are given for each metric. Each row corresponds to an experiment varying the length of the audio clip used to create spectrograms fed to the automated individual recognition feature extractor.

| Clip Duration (s) | Accuracy | ARI | NMI | FMI | V-Measure |
|---:|---:|---:|---:|---:|---:|
| 1 | 0.82 (0.71-0.86) | 0.86 (0.75-0.90) | 0.90 (0.85-0.92) | 0.86 (0.72-0.90) | 0.90 (0.85-0.92) |
| 2 | 0.85 (0.81-0.91) | 0.89 (0.85-0.93) | 0.92 (0.90-0.95) | 0.89 (0.84-0.93) | 0.92 (0.90-0.95) |
| 3 | 0.87 (0.83-0.89) | 0.93 (0.91-0.94) | 0.93 (0.91-0.95) | 0.91 (0.87-0.94) | 0.93 (0.91-0.95) |
| 4 | 0.83 (0.76-0.89) | 0.91 (0.84-0.93) | 0.91 (0.88-0.94) | 0.88 (0.82-0.93) | 0.91 (0.88-0.94) |

Table S6: Validation set performance for preprocessing experiments

Mean and range of automated recognition performance across 5 training runs are given for each metric. Rows show performance for baseline preprocessing, addition of overlay augmentation (mix-up with non-focal species sample), and addition of noise reduction preprocessing

| Preprocessing Strategy | Accuracy | ARI | NMI | FMI | V-Measure |
|---:|---:|---:|---:|---:|---:|
| baseline | 0.87 (0.83-0.89) | 0.93 (0.91-0.94) | 0.93 (0.91-0.95) | 0.91 (0.87-0.94) | 0.93 (0.91-0.95) |
| overlay | 0.90 (0.88-0.93) | 0.94 (0.93-0.95) | 0.95 (0.94-0.96) | 0.94 (0.93-0.96) | 0.95 (0.94-0.96) |
| reduce noise | 0.90 (0.86-0.93) | 0.94 (0.89-0.96) | 0.95 (0.93-0.97) | 0.94 (0.90-0.96) | 0.95 (0.93-0.97) |

Table S7: Comparison of dimensionality reduction techniques. Values show average performance across 30 runs on the validation set.

| Reduction algorithm | Output dimensions | accuracy | ARI | NMI | FMI | V Measure |
|---|---|---|---|---|---|---|
| None | 512 | 0.75 | 0.71 | 0.88 | 0.75 | 0.88 |
| t-SNE | 2 | 0.89 | 0.94 | 0.94 | 0.94 | 0.94 |
| | 3 | 0.89 | 0.93 | 0.94 | 0.94 | 0.94 |
| UMAP | 2 | 0.79 | 0.88 | 0.90 | 0.88 | 0.90 |
| | 3 | 0.84 | 0.92 | 0.92 | 0.92 | 0.92 |
| | 5 | 0.85 | 0.93 | 0.93 | 0.93 | 0.93 |
| | 10 | 0.87 | 0.93 | 0.93 | 0.93 | 0.93 |
| | 20 | 0.87 | 0.93 | 0.94 | 0.93 | 0.94 |
| | 30 | 0.89 | 0.94 | 0.94 | 0.94 | 0.94 |

Table S8: Individual identification performance on validation set when using our Ovenbird feature extractor and pre-trained baseline feature extractors. Mean and range across 30 runs are given.

| Model | Accuracy | ARI | NMI | FMI | V Measure |
|---|---|---|---|---|---|
| Ovenbird | 0.93 (0.92-0.94) | 0.95 (0.94-0.95) | 0.97 (0.96-0.97) | 0.95 (0.95-0.96) | 0.97 (0.96-0.97) |
| BirdNET | 0.49 (0.11-0.53) | 0.21 (0.00-0.28) | 0.60 (0.02-0.64) | 0.30 (0.24-0.35) | 0.60 (0.02-0.64) |
| ResNet18 ImageNet | 0.13 (0.13-0.13) | 0.01 (0.00-0.01) | 0.08 (0.06-0.08) | 0.24 (0.24-0.24) | 0.08 (0.06-0.08) |
| HawkEars | 0.13 (0.10-0.21) | 0.01 (0.00-0.03) | 0.07 (0.01-0.29) | 0.22 (0.15-0.24) | 0.07 (0.01-0.29) |

Table S9: Summary statistics of parameter posterior for the basic CJS with all individuals

| | mean | sd | 2.50% | 25% | 50% | 75% | 97.50% |
|---|---|---|---|---|---|---|---|
| phi_intercept | 0.70 | 0.02 | 0.66 | 0.68 | 0.70 | 0.71 | 0.74 |
| p_intercept | 0.89 | 0.02 | 0.85 | 0.88 | 0.89 | 0.91 | 0.93 |
| deviance | 704.65 | 1.94 | 702.72 | 703.24 | 704.05 | 705.44 | 709.75 |

Table S10: Summary statistics of parameter posterior for the full CJS model, including all individuals

| | mean | sd | 0.025 | 0.250 | 0.500 | 0.750 | 0.975 |
|---|---|---|---|---|---|---|---|
| phi_year[1] | 0.70 | 0.05 | 0.60 | 0.67 | 0.70 | 0.73 | 0.79 |
| phi_year[2] | 0.67 | 0.05 | 0.57 | 0.64 | 0.67 | 0.71 | 0.77 |
| phi_year[3] | 0.65 | 0.05 | 0.55 | 0.62 | 0.65 | 0.69 | 0.76 |
| p_intercept | 0.89 | 0.02 | 0.84 | 0.87 | 0.89 | 0.91 | 0.93 |
| sd.lphi.site | 0.20 | 0.14 | 0.01 | 0.09 | 0.18 | 0.28 | 0.53 |
| beta_slope | 0.13 | 0.15 | -0.17 | 0.03 | 0.13 | 0.23 | 0.43 |
| beta_ba | -0.03 | 0.10 | -0.23 | -0.09 | -0.03 | 0.04 | 0.18 |
| beta_canopy | 0.07 | 0.11 | -0.15 | 0.00 | 0.07 | 0.14 | 0.29 |
| phi_ohiopyle_2021 | 0.70 | 0.05 | 0.60 | 0.67 | 0.70 | 0.73 | 0.79 |
| phi_sgl034_2021 | 0.74 | 0.04 | 0.66 | 0.72 | 0.75 | 0.77 | 0.82 |
| deviance | 705.08 | 7.16 | 687.92 | 701.68 | 705.82 | 709.38 | 717.86 |

Table S11: comparison of generalized linear models for abundance with and without each point-level covariate. Blank cells indicate the variable was not included in the model. Intercept is the log of the Poisson rate lambda, the mean number of individuals per point. Covariates were standardized ("scaled") to have a mean of 0 and standard deviation of 1. Model weight is calculated by the MumIn R package.

| Intercept | Basal Area (scaled) | Canopy (scaled) | Slope (scaled) | df | Log likelihood | AICc | delta AICc | weight |
|---|---|---|---|---|---|---|---|---|
| 0.0417 | 0.146 | 0.123 | -0.091 | 8 | -712.6 | 1441.4 | 0.0 | 0.32 |
| -0.0157 | 0.145 | 0.121 | | 7 | -714.1 | 1442.4 | 1.0 | 0.19 |
| 0.0898 | 0.171 | | -0.087 | 7 | -716.8 | 1447.8 | 6.4 | 0.01 |
| 0.0340 | 0.170 | | | 6 | -718.2 | 1448.5 | 7.1 | 0.01 |
| 0.0550 | | 0.162 | -0.086 | 7 | -720.6 | 1455.4 | 14.0 | 0.00 |
| -0.0002 | | 0.161 | | 6 | -722.0 | 1456.1 | 14.7 | 0.00 |
| 0.1272 | | | -0.082 | 6 | -728.4 | 1469.0 | 27.6 | 0.00 |
| 0.0742 | | | | 5 | -729.7 | 1469.5 | 28.1 | 0.00 |

Table S12: Average number of individual male Ovenbirds detected per point across study year and block

| | 2021 | 2022 | 2023 | 2024 |
|---|---|---|---|---|
| Ohiopyle | 0.97 | 1.22 | 1.32 | 1.15 |
| SGL034 | 2.07 | 2.51 | 2.03 | 1.84 |

Table S13: Ovenbird abundance parameter estimates from the full model . Note the model uses a log link, e.g., the Poisson rate of individuals per point at Ohiopyle in 2021 is exp(0.04)=1.04. Parameters that differ from zero with $p<0.05$ are shown in bold.

| | Estimate | Std. Error | z value | Pr(>\|z\|) |
|---|---|---|---|---|
| Intercept (2021, Ohioyple) | 0.04 | 0.09 | 0.44 | 0.66 |
| **BlockSGL034** | **0.62** | **0.11** | **5.80** | **6.5e-09** |
| **2022** | **0.20** | **0.10** | **2.09** | **0.04** |
| 2023 | 0.11 | 0.10 | 1.05 | 0.29 |
| 2024 | -0.01 | 0.10 | -0.10 | 0.92 |
| Slope (scaled) | -0.09 | 0.05 | -1.73 | 0.08 |
| % **Canopy Cover (scaled)** | **0.12** | **0.04** | **2.75** | **0.01** |
| **Basal Area (scaled)** | **0.15** | **0.04** | **4.00** | **6.3e-05** |

Table S14: Summary statistics of parameter posterior for the null CJS model with all individuals with survival varying by year and by point

| | mean | sd | 2.50% | 25% | 50% | 75% | 97.50 % |
|---|---|---|---|---|---|---|---|
| phi_year[1] | 0.72 | 0.04 | 0.63 | 0.69 | 0.72 | 0.75 | 0.80 |
| phi_year[2] | 0.70 | 0.04 | 0.61 | 0.67 | 0.70 | 0.73 | 0.78 |
| phi_year[3] | 0.68 | 0.05 | 0.59 | 0.65 | 0.68 | 0.71 | 0.77 |
| p_intercept | 0.89 | 0.02 | 0.84 | 0.88 | 0.89 | 0.91 | 0.93 |
| sd.lphi.site | 0.15 | 0.13 | 0.00 | 0.03 | 0.12 | 0.24 | 0.44 |
| phi_ohiopyle_2021 | 0.72 | 0.04 | 0.63 | 0.69 | 0.72 | 0.75 | 0.80 |
| phi_sgl034_2021 | 0.72 | 0.04 | 0.64 | 0.69 | 0.72 | 0.74 | 0.79 |
| deviance | 704.31 | 5.72 | 690.09 | 701.94 | 704.86 | 707.69 | 714.29 |

Table S15: Summary statistics of parameter posterior for the residents-only CJS full model with point-level covariates on survival

| | mean | sd | 2.50% | 25% | 50% | 75% | 97.50% |
|---|---|---|---|---|---|---|---|
| phi_year[1] | 0.62 | 0.06 | 0.49 | 0.58 | 0.62 | 0.66 | 0.73 |
| phi_year[2] | 0.59 | 0.06 | 0.47 | 0.55 | 0.59 | 0.63 | 0.70 |
| phi_year[3] | 0.61 | 0.06 | 0.48 | 0.57 | 0.61 | 0.66 | 0.73 |
| p_intercept | 0.85 | 0.03 | 0.78 | 0.83 | 0.85 | 0.88 | 0.91 |
| sd.lphi.site | 0.31 | 0.20 | 0.01 | 0.15 | 0.28 | 0.44 | 0.73 |
| beta_slope | 0.04 | 0.16 | -0.26 | -0.07 | 0.04 | 0.14 | 0.35 |
| beta_ba | 0.08 | 0.11 | -0.14 | 0.00 | 0.08 | 0.15 | 0.30 |
| beta_canopy | 0.04 | 0.13 | -0.22 | -0.04 | 0.04 | 0.12 | 0.29 |
| phi_ohiopyle_2021 | 0.62 | 0.06 | 0.49 | 0.58 | 0.62 | 0.66 | 0.73 |
| phi_sgl034_2021 | 0.63 | 0.05 | 0.52 | 0.60 | 0.64 | 0.67 | 0.74 |
| deviance | 630.81 | 11.04 | 605.88 | 624.35 | 633.25 | 638.41 | 648.01 |

Table S16: Summary statistics of parameter posterior for the residents-only null model with survival varying by year and by point

| | **mean** | **sd** | **2.50%** | **25%** | **50%** | **75%** | **97.50%** |
|---|---|---|---|---|---|---|---|
| phi_year[1] | 0.63 | 0.05 | 0.52 | 0.60 | 0.63 | 0.67 | 0.73 |
| phi_year[2] | 0.61 | 0.05 | 0.51 | 0.57 | 0.61 | 0.64 | 0.70 |
| phi_year[3] | 0.63 | 0.06 | 0.52 | 0.59 | 0.63 | 0.66 | 0.74 |
| p_intercept | 0.85 | 0.03 | 0.78 | 0.83 | 0.86 | 0.87 | 0.91 |
| sd.lphi.site | 0.25 | 0.17 | 0.01 | 0.11 | 0.22 | 0.36 | 0.64 |
| phi_ohiopyle_2021 | 0.63 | 0.05 | 0.52 | 0.60 | 0.63 | 0.67 | 0.73 |
| phi_sgl034_2021 | 0.62 | 0.05 | 0.52 | 0.59 | 0.62 | 0.65 | 0.71 |
| deviance | 631.12 | 9.45 | 607.51 | 626.53 | 633.45 | 637.31 | 644.88 |

Table S17: Summary statistics of parameter posterior for the residents-only basic CJS

| | **mean** | **sd** | **2.50%** | **25%** | **50%** | **75%** | **97.50%** |
|---|---|---|---|---|---|---|---|
| phi_intercept | 0.61 | 0.02 | 0.56 | 0.59 | 0.61 | 0.63 | 0.66 |
| p_intercept | 0.86 | 0.03 | 0.79 | 0.84 | 0.86 | 0.88 | 0.91 |
| deviance | 635.76 | 2.12 | 633.77 | 634.31 | 635.11 | 636.52 | 641.35 |

#### SI I: Review of existing methods for automated individual vocal recognition

Knight et al.[1] provide an extensive review of acoustic individual identification. Here, we highlight approaches that have achieved accurate automated acoustic individual identification and review studies that performed identification of individuals not included in the training data.

Automated individual vocal recognition has been accomplished for various taxa in "closed set" scenarios where examples of each individual to be categorized are provided as training data. Historically, most individual identification approaches manually or semi-automatically selected vocalizations for analysis and measured a predetermined set of acoustic features of each vocalization, then trained supervised classification models such as discriminant function analysis or classification trees on the extracted featurest[2]. More recently, deep learning models such as convolutional neural networks have achieved high accuracy on individual acoustic classification tasks on a familiar set of individuals [3,4]. Automated visual individual recognition has also been accomplished for animal imagery for a known set of individuals[5].

Some recent studies have developed methods for automated individual category discovery, in which vocalizations from individuals not included in training data are categorized. Zhao et al.[6] developed a machine learning model to extract discriminative acoustic features from Crested Ibis calls and clustered them using UMAP for dimensionality reduction and HDBSCAN for clustering. They classified 10 individuals with 86% accuracy from focal recordings selected for high signal-to-noise ratios. Nagy and Rockwell[7] used manual feature extraction and an agglomerative clustering method to identify Eastern Screech-Owls (*Megascops asio)*, achieving 84% accuracy on 17 individuals. Sadhukhan et al.[8] similarly used manual feature extraction and clustering to identify a novel set of 20 individual Indian wolves (*Canis lupus pallipes*) with 75% accuracy. Merchan et al.[9] differentiated 10 individual manatees by clustering eigenvectors of spectrograms, though metrics of individual-level accuracy are not reported. Jensen et al.[10] perform automated acoustic feature extraction and clustering of American alligator (*Alligator mississippiensis*) bellows using focal recordings. They report a cluster purity of 61% but do not report accuracy or cluster completeness.

Additionally, in previous experiments the authors have observed that embeddings from BirdNET effectively clustered song variants of Song Sparrows (*Melospiza melodia*) from passive acoustic monitoring recordings, such that one cluster contained songs from only one individual, while multiple clusters corresponded to one individual[11].

#### SI II: Study Species

We use the wood-warbler (*Parulidae*) species Ovenbird (*Seiurus aurocapilla*) as a model species because individuals of this species produce an individually distinctive song that is stable across years and can be differentiated by inspecting the spectrograms of song recordings[12–14] (Figures 1, S1). During the breeding season (May-July), Ovenbirds sing their primary song[13,14] to defend a territory[15]. Though Ovenbirds occasionally also sing other song types, which have a completely different acoustic structure and behavioral context[14,16], we focus on the primary song type which is sung frequently in the morning[17] and has been the focus of previous work demonstrating individual acoustic signatures. Ovenbird territories (i.e., 95% use area) range from 0.1 to 1.8 hectares[15,18–20]. There is typically a buffer zone between territories[15], with singing, foraging, and nesting occurring within the territory boundary[18], though Mazerolle and Hobson[20] report that neighboring individuals forage in overlapping areas. A fraction of the males in a population (e.g., 18-37%[21]) are likely to be floaters: individuals who fail to acquire a territory and instead move through the territories of other males.

Male Ovenbirds exhibit high site fidelity across years[18,22–24]. For example, Bernard et al.[22] found that the return rate of birds to a study area in Eastern Pennsylvania was 71% and that territory centers moved by an average of 65.5 meters across their lifetime, with territories gradually drifting from the original position over a series of years. Upon returning from their first spring migration to the breeding grounds, Ovenbirds typically return to the general area where they were born but establish a territory several hundred meters from their natal home range[18,25]. In a study tracking 51 juvenile Ovenbirds, Vitz and Rodewald[25] reported a mean post-fledging dispersal distance of 1,314 m from the natal home range, with all juvenile birds moving at least 350 m from the center of the natal home range. A minority of individuals disperse large distances (e.g., tens to hundreds of kilometers[26]).

#### SI III: Comparing the labels of two reviewers

After two annotators labeled each song, we compared discrepancies between the annotators' labels using majority voting to align clusters. Then, the annotators resolved label conflicts by reviewing and discussing conflicting labels. After annotating song variants, we plotted the spatial positions of the songs colored by the song variant annotation. Because male Ovenbirds establish and sing from non-overlapping spatial territories[15], we expected that song variants would cluster spatially on the localization grids and that each spatial cluster would largely contain only one song variant. Finding multiple song variants within a spatial cluster could indicate that one bird may have produced multiple song variants or that multiple individuals used the same area within the 1-5 day period sampled. Likewise, the separation of a song variant across distant spatial clusters could indicate that multiple birds sang indistinguishable songs that were labeled as a single song variant.

#### SI IV: Automatic check of cross-correlation alignment for localized singing events

We filtered out poorly localized singing events using a novel post-processing procedure. Occasionally, the automated acoustic localization process does not correctly estimate the relative time of arrival of the Ovenbird song at each microphone, resulting in a spurious position estimate. This can occur, for instance, if a non-target sound is louder than the target sound (Ovenbird song), such that the peak cross-correlation value aligns the non-target sound rather than the target sound. If we use the estimated arrival times of the song to extract an audio clip from each microphone, the target sound will temporally align across all of the clips only if the estimated arrival times are correct. By generating a linear-valued spectrogram for each clip, then taking a pixel-wise minimum across all spectrograms, we generate a numeric array, which we term the 'min-spec', in which only sounds temporally aligned across all microphones are retained. Thus, inspecting the min-spec for the presence of the target sound (either manually or automatically) provides a rapid means of evaluating whether the target sound was properly localized. For instance, if a non-target sound appears in the min-spec and the target sound does not, the non-target sound was erroneously localized instead of the target sound. Inclusion of recorders far from the sound source can undermine this strategy, as the target sound will be quiet in the spectrogram and will erase the target sound on the min-spec. Therefore, we use only recorders within 35 m of the sound source position to create the min-spec. We normalize the min-spec peak level to the highest peak level of any of the original clips, then back-transform the spectrogram to an audio signal using the Griffin-Lim phase estimation algorithm implementation in the librosa Python package[68]. Finally, we re-run the automated Ovenbird detection process on the audio clips generated from min-specs and discard singing events if the clip's logit score for the Ovenbird class is less than -2.5, because preliminary experimentation revealed that this threshold effectively separated minspecs for which the relative arrival sounds were properly and improperly estimated.

#### SI V: Searching for individuals that occur at multiple points

In the set of Ovenbird songs used in the case study, we searched for occurrences of a single individual at neighboring sampling locations (points). For each of the 126 points, we identified the closest neighboring point. For each pair, we repeated the clustering step for the Ovenbird songs used in the case study analysis, simultaneously clustering songs from the two points. For any clusters that contained songs from both points, we inspected 10 randomly selected songs (or all songs if fewer than 10 were in the cluster) from each point to determine if the same individual was detected at both points.

In total, we found six individuals that were detected frequently at one point and in exactly one 3-second clip from the nearest neighboring point, and 3 individuals that were detected numerous times at two neighboring points. For these three individuals, the spacing between the points was 494 m (detected at one point in 2022 and the other in 2024), 627 m (detected at one point in 2021 and the other in 2023), and 324m (detected on both points in 2023).

#### SI VI: Training a feature extractor with self-supervised contrastive learning

We developed a self-supervised contrastive learning approach to training a feature extractor for acoustic individual identification. Our approach adopts machine learning techniques that are well-suited to the problem of individual identification, in which the model must generalize to new individuals beyond the training data. Unlike supervised classification, where the model's task is to predict the correct class labels of a sample, in contrastive learning, the task is to represent samples in a feature space such that similar samples are close together and dissimilar samples are far apart. Furthermore, our approach is self-supervised, meaning it does not rely on human annotation of individual songs for the training data. For tasks that involve clustering categories (in our case, individual birds' songs) not present in the training data, self-supervised machine learning approaches can outperform supervised approaches because they learn to extract and discriminate general features in the data, rather than only the features relevant to a specific set of classes.

Our model architecture uses a CNN backbone and replaces the classification layer with a projection head. We start with a ResNet18 CNN model architecture trained on ImageNet to classify images. We remove the final fully connected classification layer and add a two-layer projection head similar to that used in other contrastive learning approaches[27]. We also modify the first convolutional layer to accept one-channel inputs.

Model training hyperparameters, preprocessing, and augmentation are the same as for supervised classification training (see main text). Scripts in the GitHub repository provide Python implementations of the model (model.py) and loss (loss.py).

Projection head structure:

| Layer | Output shape |
|---|---|
| (feature extractor backbone) | (batch_size, 512) |
| Batch Normalization | (batch_size, 512) |
| ReLU activation | (batch_size, 256) |
| Linear | (batch_size, 256) |
| Batch Normalization | (batch_size, 128) |

At a high level, we train the feature extractor using at least two types of incentives:

(1) Songs recorded at distinct sampling points do not contain the same bird, and therefore should have dissimilar feature vectors

(2) Two copies of the same song clip, with different augmentations applied, should have similar feature vectors

We optionally include each of two additional incentives:

(3) Songs recorded at the same sampling point should have more similar feature vectors

(4) Pseudo-label similarity: We intermittently use feature vectors to cluster songs, then treat these clusters as pseudo-labels. Songs with the same pseudo-label should have similar feature vectors.

We formulate each of these incentives as a log-sum-exponent of pairwise feature similarity, and use them to construct a four-part contrastive loss. Following the SimCLR contrastive learning approach[27], the loss function is evaluated on the outputs of the projection head. We define a composite contrastive loss with four components:

loss = different_point_loss + same_clip_loss + + same_psuedolabel_loss

We first calculate the pairwise similarity (dot product) of each sample's feature vector as

similarity_matrix = (features * features.T) / temperature

Where features is a matrix containing the 1x512-dimensional feature vector created by the backbone for all samples in a batch. We set temperature=0.1 to increase the magnitude of the similarity values. The first two loss terms are always included.

1. First, the different_point_loss term penalizes feature similarity for pairs of samples from different survey points, since we assume these samples come from different individuals:

   different_point_loss = mean(log(sum(exp(similarity_matrix)*different_point_mask) + eps))

   Where different_point_mask contains 0s and 1s, such that only pairs of samples that are not from the same point contribute to the loss term, and eps is a small value for numerical stability.

2. The same_clip_loss term incentivises feature similarity for duplicated versions of the same audio clip that are stochastically augmented multiple times. The term computes the average log-sum-exponent of the pairwise similarity of all pairs of samples from the same original clip.

   same_clip_loss = - mean(log(sum(exp(similarity_matrix)*same_clip_mask) + eps)

   Where same_clip_mask contains 0s and 1s, such that only pairs of samples with the same singing event contribute to the loss term.

3. We optionally include a same_point_loss term to incentivize feature similarity for pairs of samples from the same survey points (the samples may contain the songs of the same or different individuals):

   same_point_loss = mean(log(sum(exp(similarity_matrix)*same_point_mask) + eps))

   Where same_point_mask contains 0s and 1s, such that only pairs of samples that are not from the same point contribute to the loss term.

4. We optionally include the same_pseudolabel_loss term, which incentivises feature similarity between pairs of samples that have the same pseudo-label. Pseudo-labels are generated by clustering training set samples on a point-by-point basis at the end of each training epoch. The loss term averages over all pairwise comparisons of samples from the same pseudo-label for a training batch. Samples not included in any clusters during stage 2 are ignored by this loss. The loss term is computed as a log-sum-exponent over relevant pairwise comparisons,

   same_pseudolabel_loss = -mean(log(sum[exp(similarity_matrix)*same_pseudolabel_mask] + eps))

   Where same_pseudolabel_mask contains 0s and 1s, such that only pairs of samples with the same pseudo-label contribute to the loss term.

When using the self-supervised learning approach (i.e., including term 4 of the loss function), model fitting proceeds by iterating through (Stage 1) pseudo-labeling the acoustically localized songs and (Stage 2) training one epoch with contrastive learning.

#### SI VII: Detection, localization, and annotation of Ovenbird songs from localization grid data

The automated Ovenbird detector produced 808,639 detections of Ovenbird vocalizations across all 13 localization arrays (15,675-210,029 per array). After filtering with the min-spec procedure (see Methods), the automated localization procedure produced 60,961 localized Ovenbird singing events (169-12,084 per array). Our manual review process produced 3,963 audio clips from 717 singing events. From these singing events, we annotated 45 individual Ovenbird song variants, with 1-8 song variants per localization array. We provide a public dataset containing the spatial information, individual labels, and associated audio clips for these Ovenbird songs (Environmental Data Initiative edi.2049.1), and an interactive web app for exploring audio samples and maps of individuals' songs (https://ovenbird-id.streamlit.app/).

Annotator agreement for individual song labels was 98.8%. Most label disagreements came from accidental typing errors during the annotation process. There were two instances in which one annotator noticed acoustic details differentiating two song variants that the other annotator had missed (examples of these songs are provided in Figures S16 and S17). Plotting annotated songs onto the localization grids revealed that, as we hypothesized, individual Ovenbird song variants tended to align with spatial clusters, which we interpret as individual males' territories (Figure 3). This finding supports the findings from previous work that each male Ovenbird sings a single song type that is self-consistent and distinguishable from nearby males[13,14].

#### SI VIII: Individual identification feature extractor training experiment results

With the supervised classification approach, cross-entropy loss and binary cross-entropy loss performed best on the validation set (mean accuracy of 0.87 across 5 runs). ArcFace loss also performed well, achieving its best performance with two subcenters per class (mean accuracy 0.82). All variations of the contrastive learning approach performed poorly, with a mean accuracy of 0.12 or lower. We provide a Weights and Biases report (https://bit.ly/ovenbird-training-experiments) for detailed investigation of experimental results. We also provide a table containing detailed parametrization and metrics for each training run on the GitHub repository (training_experiment_results.csv).

The best-performing backbone was the default and smallest, ResNet 18. Resnet-50 had marginally worse performance, and HawkEars had substantially worse performance. This result could be a product of the smaller embedding output size of ResNet 18 (512, compared to 1024 for ResNet-50 and 2048 for HawkEars) being more suitable for the downstream clustering task, or a result of the smaller number of trainable parameters in ResNet 18.

Performance increased with the size of the training set and with the number of unique acoustic sampling locations included in the training set (Figure 4, Table S2, S3). Most of the performance improvement was gained by reaching 10,000 training samples, with small additional improvements for 50,000 and all 94,378 clips (Figure S4a). Likewise, performance was moderate when only 16 sampling locations were included in the training set (69% validation set accuracy), excellent with 64 locations (85%), and marginally better with 128 or 234 locations (87%, Figure S4b).

Noise-reduction and overlay augmentation (mix-up of Ovenbird song clip with a non-Ovenbird clip) each improved model performance from 87% to 90%. The best-performing input clip length was 3 seconds, with other clip lengths performing marginally worse (1s: 82%, 2s: 85%, 3s: 87%, 4s: 83%). Based on these experiments, we trained a full model on the entire training set using a ResNet18 backbone with 3-second clip inputs and included noise-reduction and overlay in preprocessing.

#### SI IX: Dimensionality reduction experiment results

Clustering accuracy was much higher when the feature vectors were dimensionally reduced using t-SNE or UMAP (validation accuracy 89%) than without dimensionality reduction (75%, Table S7). Both t-SNE and UMAP performed well, but UMAP only matched the performance of t-SNE when the number of output dimensions was increased to 30. Comparing UMAP with 30 output dimensions to t-SNE with 3 output dimensions, UMAP produced much less consistent performance (accuracy standard deviation across runs 2.0% compared to 0.5% for t-SNE). Because saving 3-dimensional embeddings requires less storage and results in faster clustering, and because it offered more consistent performance, we used t-SNE with 3 output dimensions for the case study.

#### SI X: Evaluation of automated species classification performance

##### Evaluating species classifier accuracy

By annotating 1000 randomly selected 3-second audio clips from the Localization Dataset, we created a species classifier evaluation set containing 90 positives containing Ovenbird song and 896 negatives, and discarded 14 clips where the reviewer was uncertain or where Ovenbird song only occurred in less than 0.3 seconds at the beginning or end of the clip. After applying the HawkEars species classification model to this test set, we reviewed clips labeled as negative but receiving a logit score over 0, and found that two of three clips were incorrectly annotated. After adjusting these labels, the Hawkears model achieved excellent performance for Ovenbird song binary classification (AU-ROC: 0.985, AP: 0.926). This annotated dataset is available under an open-source license (Dryad: in prep).

##### Realized classifier precision and recall in passive acoustic monitoring datasets

At the logit-score threshold of -1.0 used for detecting vocalizations on the Localization Dataset, precision was 0.91 and recall was 0.83. At the threshold of 1.0 used for the Longitudinal Dataset case study, precision was 0.97, and recall was 0.69. When removing Ovenbird clips where other species were detected at a threshold of 0.0, precision was 0.96, and recall was 0.60. When muting all but the central 1-second of the clip before HawkEars classification, the threshold score of 0.0 used in the case study resulted in a precision of 1.0 and a recall of 0.17. Low recall probably results from the average-pooling layer of the machine learning classifier, and other approaches to finding centered songs deserve attention in future work.

**Filtering automated detections before individual identification: challenges and future directions**

When performing end-to-end automated individual identification on acoustic monitoring data, we found it critical to filter the detections produced by an automated detector such that only centered, unobscured songs were included as inputs to clustering. Regardless of which embedding model was used, if all Ovenbird detections from the automated classifier were used in the individual identification procedure, the embeddings formed a single large cluster in which individuals could not be distinguished. We chose to implement simple heuristics for filtering detections to those of high quality, such as removing audio clips where other species were detected and re-running the classification algorithm with the beginning and end of the clip muted so that only songs occupying the center of the clip were detected. However, the muting strategy lowered recall from 0.60 to 0.17. We experimented with alternatives for locating and isolating foreground sounds, but found that deep-learning-based sound source separation (MixIt[28]) was ineffective, while gradient backpropagation (GradCAM[29]) was effective but prohibitively computationally expensive. In the future, alternative approaches to filtering detections and isolating foreground sounds could improve the pipeline's performance. The development of a targeted source separation algorithm that separates the sound of a specific class (e.g., species song) from other sounds would be particularly valuable, as it would convert noisy recordings from passive acoustic monitoring data into clean focal recordings where irrelevant sounds have been removed.

#### SI XI: Results and discussion of survival and abundance analyses

The automated Ovenbird detection procedure produced 147,879 3-second Ovenbird audio clips across the 126 study points and 4 years (Figure S14). Our automated individual identification process produced 470 song clusters across the 126 points (including data from all 4 years). Points (N=126) had 0 to 10 clusters each (median 4, mean 4.01), and 117/126 points had at least one cluster. After removing contaminated clusters and merging clusters containing the same song variant (Phase 1 review), 405 clusters remained. Points then had 0 to 7 clusters each (median 3, mean 3.46). During this review process, 64 clusters were removed because songs were distant and unidentifiable at the individual level; 18 clusters were removed because no single song variant was dominant; 7 clusters were removed because they did not contain Ovenbird songs; and 72 clusters were merged with an existing cluster containing the same song variant. From retained clusters, 12% of reviewed samples were removed because they either were too distant to confidently identify or did not match the dominant song variant of the cluster.

We fit Cormack-Jolly-Seber (CJS) models to the four-year detection histories for all individuals and only resident individuals (those detected on at least 4 of 12 days in a given year). The basic CJS model, including all individuals, found that apparent survival was 0.70 (95% credible interval 0.66-0.74) and annual detection probability was high (0.89, 95% CI 0.85-0.93, posterior distribution and summary statistics in Figure S7 and Table S9). Parameter estimates were similar for the null model (Table S14). The residents-only full model found that apparent survival of residents was lower (0.61, 95% CI 0.56-0.66, Figure S7, Table S15), which is expected since this approach removes detections of individuals detected on fewer than 4 days, and apparent survival estimated by this model thus represents the probability of both returning and establishing residency. Survival and recapture probability estimates from the base and null models for only residents were similar to those from the full model (Tables S16, S17).

Because apparent survival does not distinguish between emigration and death, and Ovenbirds' may typically shift territory centers by an average of 66 meters per year[22], the spatial and temporal extent of resighting surveys could correlate with modeled apparent survival even when accounting for imperfect detection. Specifically, increasing the surveyed area will convert some emigrated birds into returning birds, raising apparent survival. Increasing the temporal extent of the survey could have the same effect if birds slightly outside of the surveyed area occasionally make forays into the surveyed area. Acoustic resighting through passive acoustic monitoring, which achieves high temporal sampling coverage, may therefore produce higher estimates of apparent survival than traditional resighting methods, though this will depend on the area sampled by each method.

Our full model estimated that apparent survival was marginally highest in the first to second year, and point-level apparent survival was not explained by habitat covariates (Table S10). In the full model, survival was estimated to be 0.70 in years 1 to 2 (95% credible interval: .60-.79), 0.67 (.57-.77) in year 2 to 3, and 0.65 (.55-.76) in year 3 to 4. Estimates of apparent survival with the null model that did not include habitat covariates were approximately 2% higher than with the full model. Yearly survival was consistent across years in the residents-only model (year 1-2: 0.62, year 2-3: 0.59, year 3-4: 0.61). We did not find evidence of survival varying with study area (Ohiopyle versus SGL 34) or site-level habitat covariates in models for all individuals or for residents (parameter posteriors in Figure S9, S15).

#### SI XII: Estimating abundance and survival with and without automated individual recognition

In the absence of an automated tool for individual vocal recognition, could survival and abundance be measured by selecting random subsets of the passive acoustic monitoring dataset and annotating Ovenbird songs to individuals? To answer this question, we ran simulations to investigate how random annotation of Ovenbird songs would compare to using automated individual identification. We found that randomly reviewing Ovenbird song detections at each point would discover fewer individuals and under-estimate abundance (Figure S11a,b). Random annotation also generates lower resighting rates even with high annotation effort, resulting in inconsistent and biased underestimates of apparent survival from CJS models (Figure S11c). Thus, using an automated individual recognition approach to identify individuals across the full extent of the passive acoustic monitoring dataset, rather than annotating a random sample, is critical to accurately measuring Ovenbird abundance and survival.

#### SI XIII: Do song variants correspond one-to-one with individual Ovenbirds?

Because we did not mark or capture birds in this study, we should critically consider how likely it is that Ovenbird song variants (whether clustered by humans or the automated approach) have a one-to-one correspondence with individual birds. In particular, we should be concerned with the possibilities that (1) individual Ovenbirds' songs cannot be reliably distinguished by humans or automated analyses; (2) multiple individuals are assigned to a single song variant (cluster); or (3) multiple song variants (clusters) belong to the same individual. We will address each of these possibilities in light of our findings.

First, in agreement with previous studies[13,14], we found strong evidence that individual Ovenbirds' songs can be reliably identified both by humans and automated procedures. Our human-labeled annotations of individual Ovenbirds corresponded well to spatial clusters on the localization arrays that delineate individual birds' territories (Figure 3). Between-annotator agreement was also high (98%). Furthermore, our automated approach, which was trained by learning to separate songs from a different Ovenbird population in Pennsylvania, USA, also produced very similar annotations to the two human annotators for the localization grids (94-99% accuracy). Therefore, the discriminative acoustic features present in Ovenbird songs in Pennsylvania, USA, were sufficient for distinguishing individuals in a geographically distant population in Alberta, Canada. These results indicate that the acoustic features of Ovenbird songs can be used for accurate individual identification.

Second, we found evidence that multiple individuals are unlikely to sing indistinguishable song variants, but spatial clusters of Ovenbird songs from the localization arrays, which we interpret as individual territories, typically contained only one song variant (Figure 2). In theory, individuals could have indistinguishable songs either because (a) populations with low rates of emigration and immigration can form song neighborhoods of individuals with very similar sons, or (b) in a large set of individuals, some will have indistinguishable songs by chance. Regarding a, we did not find evidence that neighboring birds have very similar songs: songs taken from single sampling locations in Pennsylvania had diverse acoustic features, as did songs from neighboring territories on the Alberta localization grids (see interactive web page: [22]). Regarding b, our approach was able to discriminate over 200 individual clusters from a pooled set of songs while retaining high purity within clusters (Figure 5b). Therefore, we infer that individual Ovenbirds typically can be distinguished from others by song.

Third, we found evidence that Ovenbirds are unlikely to sing multiple song variants. This finding is consistent with the observation from Ehnes and Foote[13] and Lein[14] that Ovenbirds sing a single, self-consistent song. Taken together, these results provide compelling evidence that Ovenbird songs largely have a one-to-one correspondence with individual birds, though we cannot rule out the possibility of rare exceptions.

Because we did not physically mark individuals, some level of uncertainty remains in acoustic recapture analysis. Misidentification from acoustic monitoring is analogous to errors that occur when resighting physically marked individuals[30] and can result in biased estimates of survival. In particular, we highlight three scenarios that could cause identification errors and potentially bias survival estimates from acoustic recapture. First, young birds could learn songs from a nearby tutor and reproduce them faithfully in a later year. Songbirds such as Ovenbird are known to learn their songs before departure from their natal home range or following the settlement of their territory the following spring, using nearby conspecifics as tutors. Males of some songbird species can produce very faithful copies of their tutors' songs[31], though this phenomenon has not been reported for Ovenbird. In the context of a multi-year survival analysis, this type of error would lead us to incorrectly believe the tutor is still surviving and returning to the site, resulting in an overestimate of survival. Second, if different individuals at a survey point sing indistinguishable songs, placing them into a single song cluster would lead to an under-estimate of abundance and potentially an overestimate of survival. Third, some individuals might sing more than 1 unique song, or change their song over time, which would lead to an overestimate of abundance and potentially to an underestimate of survival.

#### SI XIV: Opportunities and challenges related to statistical analyses of acoustic recapture data

Developing appropriate statistical methods for extracting ecological insights from individual passive acoustic recapture histories will be a complex and nuanced topic of study. There is a rich body of work on modeling survival and other parameters from individual recapture histories while accounting for phenomena such as heterogeneous detectability, transient individuals, and spatial and temporal autocorrelation[32,33]. Acoustic monitoring data are particularly well suited for robust design models where secondary sampling periods are nested within primary sampling periods[34,35]. Spatial capture-recapture models[32] provide a means of separating true annual survival from emigration, but likely require different spatial sampling designs than those currently used for passive acoustic monitoring: typical PAM study designs use spacing that minimizes the chance of capturing the same individual on multiple devices, but spatial capture recapture models rely on capturing individuals at multiple sampling locations to estimate dispersal or movement kernels. Overall, it will be necessary to confront properties of the acoustic recapture data type that differ from conventional capture-recapture protocols and may violate assumptions of existing statistical approaches, in particular those resulting from high temporal survey coverage.
